# Divergence and Remarkable Diversity of the Y Chromosome in Guppies

**DOI:** 10.1101/2020.07.13.200196

**Authors:** Pedro Almeida, Benjamin A. Sandkam, Jake Morris, Iulia Darolti, Felix Breden, Judith E. Mank

## Abstract

The guppy sex chromosomes show an extraordinary diversity in divergence across populations and closely related species. In order to understand the dynamics of the guppy Y chromosome, we used linked-read sequencing to assess Y chromosome evolution and diversity across upstream and downstream population pairs that vary in predator and food abundance in three replicate watersheds. Based on our population-specific genome assemblies, we first confirmed and extended earlier reports of two strata on the guppy sex chromosomes. Stratum I shows significant accumulation of male-specific sequence, consistent with Y divergence, and predates the colonization of Trinidad. In contrast, Stratum II shows divergence from the X, but no Y-specific sequence, and this divergence is greater in three replicate upstream populations compared to their downstream pair. Despite longstanding assumptions that sex chromosome recombination suppression is achieved through inversions, we find no evidence of inversions associated with either Stratum I or Stratum II. Instead, we observe a remarkable diversity in Y chromosome haplotypes within each population, even in the ancestral Stratum I. This diversity is likely due to gradual mechanisms of recombination suppression, which, unlike an inversion, allow for the maintenance of multiple haplotypes. In addition, we show that this Y diversity is dominated by low-frequency haplotypes segregating in the population, suggesting a link between haplotype diversity and female-preference for rare Y-linked colour variation. Our results reveal the complex interplay between recombination suppression and Y chromosome divergence at the earliest stages of sex chromosome divergence.

## Introduction

Sex chromosomes diverge from each other once recombination between the X and Y chromosome is halted ^1,2^. Despite the prevalence and repeated origin of sex chromosomes ^3^, we still know little about the initial stages of X-Y divergence. In particular, it remains unclear exactly how recombination is suppressed between emerging sex chromosomes. Classic models assume that recombination is instantly and completely arrested when one chromosome undergoes an inversion ^4^. However, empirical studies have suggested that the earliest stages of recombination suppression may be due to shifts in local recombination hotspots ^5^, which could, at least initially, provide an incomplete barrier to recombination. Moreover, inversions are rare events, and fixation of a Y-linked inversion as a mechanism to achieve recombination suppression would lead to a limited number of Y haplotypes. If recombination is curtailed through means other than an inversion, substantial initial haplotype diversity can persist within the non-recombining region. The mechanism of recombination suppression can leave fundamentally different patterns of diversity on emerging Y chromosomes.

Observations of Y-linked male colour traits in guppies (*Poecilia reticulata*) ^6–10^, helped inspire theories of sex chromosome formation ^11^, and observations that Y-linkage of colour varies by population ^6,12–14^ have fuelled speculation of a link between sex chromosome divergence and female preference for Y-linked male colour combinations ^15–18^. Curiously, there appears to be a surprising diversity of Y-linked colour haplotypes ^19,20^ which is somewhat counterintuitive as the combination of purifying selection and linkage effects on Y chromosomes typically combine to remove diversity from these regions ^21^.

Guppies have recently resurfaced as a model for the genomic study of sex chromosome formation. Recent work has suggested that *P. reticulata* shows evidence of early Y degeneration at the distal end of Chromosome 12, in a region that is also ancestral to *Poecilia wingei*, its sister species (Stratum I) ^15,17,22–24^. This non-recombining region is consistent with previous cytogenetic and linkage studies ^8,9,25–28^, and a genetic map of the sex determining region ^29^. In addition, the extent of X-Y divergence varies substantially among populations of *P. reticulata* and between *P. reticulata* and *P. wingei* ^15,22,24^. This suggests that recombination is also rare beyond Stratum I, either because recombinants are selected against in natural populations, or the rate of recombination is sufficiently low that does not fully counter the accumulation of Y-specific mutations. However, it is important to note that others have not recovered support for this region of recombination suppression using slightly different genomic approaches ^16^, and it has been suggested that the guppy Y chromosome lacks any discernible divergence from the X, despite cytogenetic work showing X-Y differentiation ^30–32^.

The guppy sex chromosome system is surprisingly old, as it is shared with *Poecilia picta* ^22^, suggesting it originated at least 20 million years ago ^33^. The age of the sex chromosome system, coupled with the fact that the sex chromosomes in *P. picta* are highly diverged and possess a mechanism of complete X chromosome dosage compensation in males ^22^, indicates that other forces are counteracting the degeneration normally associated with non-recombining regions to maintain the overall integrity of the Y in *P. reticulata*.

In the northern range mountains of Trinidad, downstream and upstream river populations are known to differ in male colour patterns and other important life-history traits due to differences in predator and food abundance ^34,35^, and these complex phenotypes have been shown to evolve rapidly and repeatedly ^36^. In order to understand sex chromosome divergence in this system, identify Y-specific sequence, and determine the mechanism of recombination suppression, we generated linked-read sequences from multiple males and females from both upstream and downstream population pairs from three rivers in Trinidad, and used these data to build high-quality population-specific genome assemblies.

Given the controversy over the extent of X-Y divergence in this species ^15–18^, we first independently replicated our previous results ^15,22^. We again recovered evidence of both a region of X-Y divergence shared among populations (Stratum I), and a convergently evolved region of greater X-Y divergence in upstream compared to downstream populations (Stratum II). We then use linked-read assemblies to show a significant accumulation of male-specific sequence and male-linked SNPs in Stratum I, and reveal an astonishing diversity in Y haplotype sequence both among and within populations. We show that this diversity is not associated with an inversion, and instead suggest that the lack of a structural mechanism of recombination suppression allows for the maintenance of large numbers of Y haplotypes. Taken together, our results reveal the initial stages of sex chromosome divergence and the evolutionary processes that allow Y diversity to be maintained.

## Results

We collected and obtained whole-genome sequencing reads for 120 wild-caught individuals, 20 male and 20 female *P. reticulata* samples for each of the Aripo, Quare and Yarra rivers in Trinidad, equally divided between downstream and upstream populations. We used a combination of 10x Genomics linked-read sequencing and paired-end Illumina sequencing (see Materials and Methods and Supplementary Table S1), and after filtering alignments to the reference genome (see Materials & Methods), we recovered an average effective coverage of ∼30X for each male and ∼20X for each female.

In order to remove potentially confounding effects of the underlying genetic variation between rivers, we constructed river-specific reference genomes using the best female linked-reads’ *de-novo* assembly, based on scaffold N50 and phase block N50 values (Supplementary Table S2). In all cases, these assemblies showed a high completeness and contiguity with N50 of >1 Mb, 3 Mb and 8 Mb for Yarra, Aripo and Quare genomes, respectively ^37^ (Supplementary Table S3). We anchored our assemblies to the published guppy genome, and identified a clear inverted segment of the first 10 Mb on the sex chromosome (Chromosome 12) (Supplementary Figure S1). This rearrangement is shared across the three rivers we collected from, suggesting that it may be a unique feature of the guppy reference genome not present in any of our natural populations. It is also not present in related outgroup species or our laboratory guppy population ^22,24^.

### Sex chromosome divergence

Degeneration or significant divergence of the Y chromosome results in reduced male coverage when mapped to a female reference genome, and therefore the ratio of male to female mapped reads can be used to identify regions where the Y chromosome has significantly degraded compared to the X ^22,38–41^. We have previously used this approach to identify a small region of significant Y divergence in all of the populations we assess here ^15^, which was present in the common ancestor with *P. wingei* ^22,24^, designated as Stratum I. Supporting these previous findings, we again find evidence for this region in all our populations at the distal end of Chromosome 12, largely due to a drop in male mapping at 21-22 Mb and 25-26 Mb (Supplementary Figures S2 and S3). Stratum I also exhibits elevated male:female F_ST_ in all our assessed populations (Figure 1). Together, these results are consistent with previous cytogenetic evidence ^25–27^ suggesting that Stratum I contains the male sex determining region (SDR), most likely within the intervals of 21-22 Mb or 25-26 Mb.

**Figure 1.**
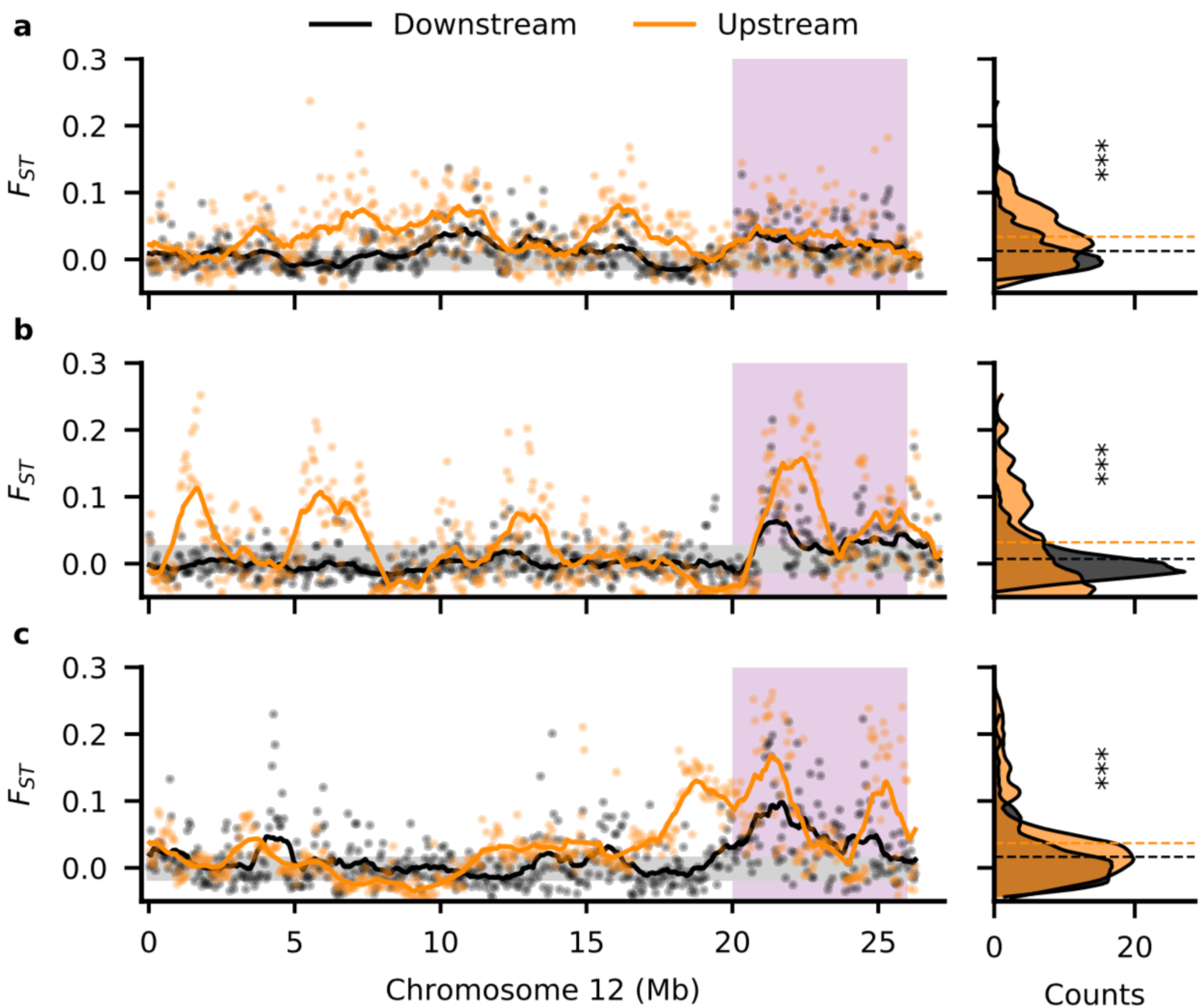
Increased divergence between sexes in the upstream populations. Male-female F_ST_ along Chromosome 12 for downstream (black) and upstream (orange) populations in the Aripo (**a**), Quare (**b**) and Yarra (**c**) rivers. Stratum I is shaded in pink. F_ST_ was calculated in non-overlapping windows of 50 kb. The 95% confidence interval, inferred from bootstrapping autosomal regions, is shaded in grey. The right-side density plots show the frequency (counts) of windows for the F_ST_ values. Dashed horizontal lines indicate the mean F_ST_ in each population. All upstream populations showed a significant increase in F_ST_ relative to the downstream populations (***, p-value < 0.001).

We previously also observed elevated male SNP density ^15^ in each of our upstream populations of *P. reticulata* across a larger proportion of the sex chromosome compared to downstream populations, which we designated as Stratum II. However, this observation was not corrected for the large inversion on this chromosome that is present in the reference genome but absent from our study populations (Supplementary Figure S1). The region of elevated male SNP density is also apparent in *P. wingei* ^22^, where it formed independently ^24^. We expect that the accumulation of male-specific mutations on the Y will lead to increased allelic difference (F_ST_) between males and females. We observe significant increases in F_ST_ in replicate upstream compared to downstream populations along the length of Chromosome 12 (*P* < 0.001, Kruskal-Wallis test; Figure 1). Once correcting for the inversion, our results suggest that Stratum II in fact encompasses a larger extent of Chromosome 12 than we previously observed.

### Male-linked polymorphisms

In a male heterogametic system, Y-linked alleles are only transmitted through male gametes. Thus, Y-linked regions that still retain homology to the X chromosome will show greater heterozygosity in males (XY) compared to females (XX). With a high number of Y haplotypes, explained below, we do not expect all Y-linked SNPs to be present in all males, and we therefore examined SNPs absent in all females but present in a subset of males. We observe an excess of such male-linked SNPs on Chromosome 12 (Supplementary Table S4), as this chromosome was the only one in the genome where the observed number of male-linked SNPs was higher than the false positive rate (obtained from random resampling in all populations, 10,000 iterations) accounting for chromosome size. Furthermore, the distribution of male-linked SNPs on Chromosome 12 was strongly skewed, with significantly more sex-linked SNPs than expected by chance (*P* < 0.001, X^2^ test) in Stratum I based on its total length and SNP content (Figure 2). Within Stratum I, we generally observe increased levels of male-linked SNPs at 21-22 Mb and 25-26 Mb, consistent with the area of greater Y divergence (Figure 1, Supplementary Figures S2 and S3).

**Figure 2.**
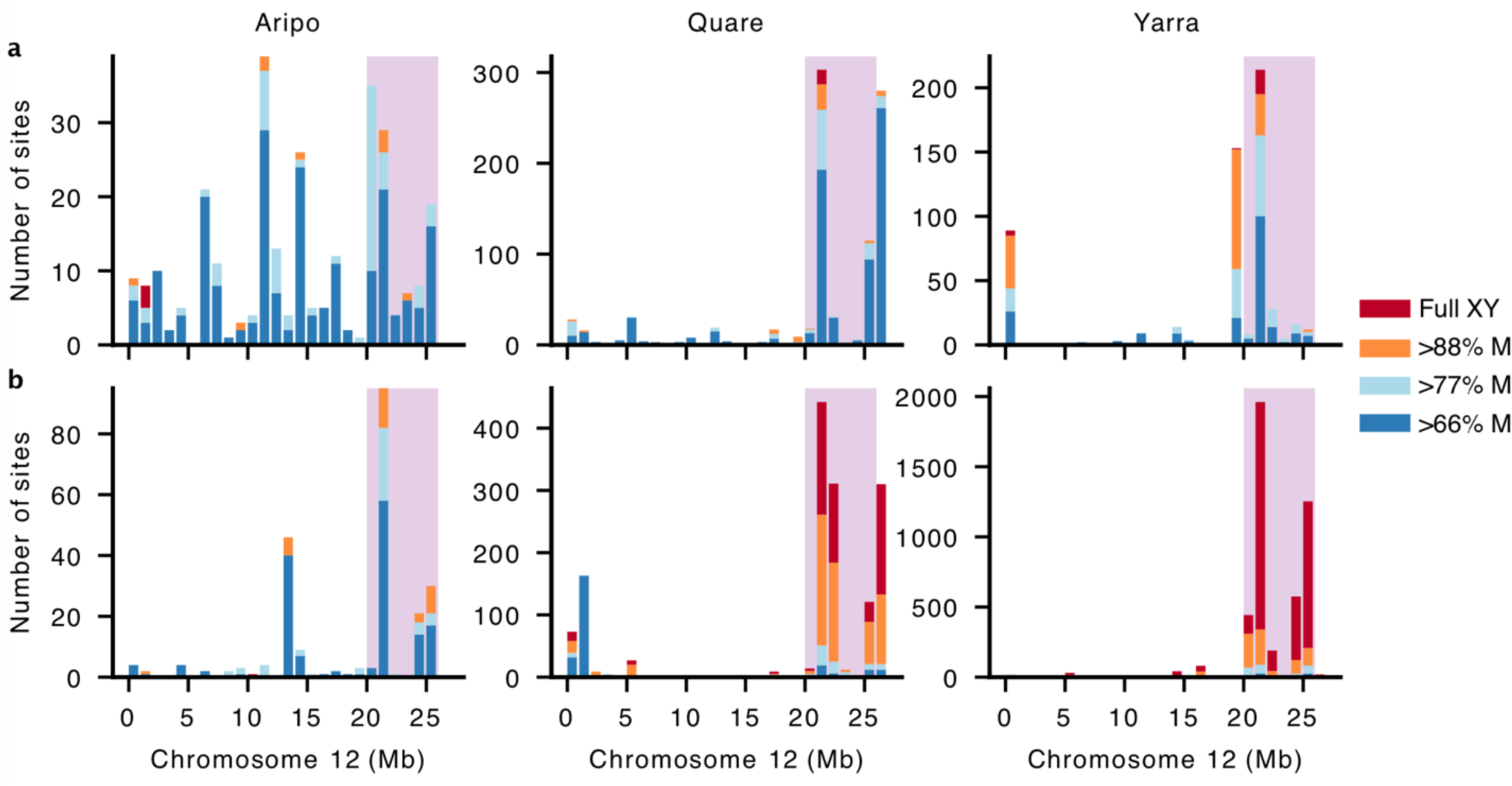
Chromosome 12 shows a skew distribution of male-linked SNPs in Stratum I, particularly for upstream populations. The number of male-linked SNPs is shown for downstream (**a**) and upstream (**b**) populations. The number of SNPs compatible with complete male-linkage (in 100% of males is shaded in red, male-linked in >88% (all but 1 male) in orange, >77% (all but 2 males) in light blue, and >66% (all but 3 males) in dark blue. Note that the scale of the Y axis differs between populations. SNPs are plotted in 1 Mb windows. Stratum I is shaded in pink.

We then analysed fully male-linked SNPs, those heterozygous in all males and homozygous in all females, on Chromosome 12 and observed marked differences across rivers. Consistent with a previous genotyping study of a downstream population from Aripo ^16^, we also failed to recover fully male-linked SNPs in Stratum I for this watershed. These findings, however, contrast markedly with the other populations. In both Quare and Yarra populations, the distribution of fully male-linked SNPs was again significantly skewed (*P* < 0.001, Χ^2^ test), with either all (Quare downstream) or most (>62% in Quare upstream and both Yarra populations) of fully male-linked SNPs present in Stratum I. Interestingly, genetic diversity on the sex chromosome and autosomes, estimated as the proportion of segregating sites (Watterson’s theta), was highest in Aripo compared to the other watersheds (Supplementary Table S5). Genetic diversity is influenced by many factors, and is known to show a strong positive correlation with variation in recombination rates ^42^, suggesting that there may be genetic variation in this watershed for higher overall recombination rates.

The total proportion of all male-linked SNPs found in Stratum I also varied extensively between populations of the same river. In particular, upstream populations showed 1.4X (Aripo), 1.9X (Quare) and 15.6X (Yarra) more male-linked SNPs in Stratum I than their corresponding downstream populations. This difference was even more striking when considering only fully male-linked SNPs, with the upstream populations showing 21.5X (Quare) and 178.8X (Yarra) more full sex-linked SNPs than the respective downstream populations (Figure 2). Altogether, our results suggest higher sex-linkage in the distal end of Chromosome 12 and a much stronger association of sex-linkage in upstream populations, which is consistent with our previous study on these guppy populations ^15^. Despite these differences, a PCA analysis of all SNPs from Stratum I does not fully result in the clustering of samples by sex (Supplementary Figure S4), suggesting incomplete lineage sorting of many Y alleles.

### Lack of inversion between X and Y chromosomes

Inversions are often implicated as a major driver of recombination suppression in sex chromosomes ^41,43,44^, and can accelerate lineage sorting by creating a strong bottleneck when a rare inversion haplotype becomes fixed in a population. Given the curious lack of lineage sorting for the Y chromosome (Supplementary Figure S4), we used the linked-read information from 10x Genomics sequencing to search for differences between males and females in the aligned distance of barcodes. This method has considerably more power than short-read sequencing to detect rearrangements because the long molecules are more likely to span the rearranged breakpoints.

If an inversion has formed between the X and Y chromosome, we would expect males to be heterozygous for this structural variation and that it would be absent in females because the reference genome is also from a female. Although we could detect distinctive barcode overlaps between distant positions of Stratum I across the three rivers, we failed to identify any consistent difference between females and males (Supplementary Figure S5). This suggests that any potential rearrangement involving Stratum I is shared between the sexes, and therefore not Y-specific (Supplementary Figure S1a). We therefore found no evidence of an inversion on Stratum I, possibly explaining the unusual Y diversity and lack of complete lineage sorting in this region.

### Sequence analysis of Y haplotypes

To further investigate the genetic diversity of the guppy Y chromosome, we used our linked-read sequencing data to computationally phase all genotypes in our populations (see Materials and Methods). We estimated a phase switching error of <5% for all linked-read samples, with the exception of one Quare male and one Yarra female (Supplementary Figure S6), indicating that this approach provides high phasing accuracy. We then used gene trees in non-overlapping windows of 100 SNPs to infer Y-linked haplotypes. Briefly, if a region of the Y chromosome is linked to the SDR and recombination suppression has led to complete lineage sorting of X and Y alleles, it will only be present in males and, therefore, will form a monophyletic clade composed exclusively of male samples.

Regions where >66% of males formed a monophyletic clade largely clustered in Stratum I (Figure 3a), again predominately between 20-22 Mb and 24-26 Mb. Although we also observed some XY topologies at low frequency in the pseudoautosomal region (PAR), these are presumably caused by an overall reduced recombination rate (heterochiasmy) in male guppies ^16^. The frequency of XY topologies in Stratum I was highest in Yarra, intermediate in Quare and lowest in Aripo (Figure 3b), in line with the distribution of sex-linked SNPs in these rivers (Figure 2). Interestingly, most of the Y haplotypes detected were shared between populations, with very few (4 regions within Stratum I of Aripo, 3 in Quare and 24 in Yarra Rivers, corresponding to 12.5%, <3% and <7% of the total number of XY topologies in this region) unique to a single population (Figure 3b). This is expected if recombination on Stratum I is ancestral to *P. reticulata* populations in Trinidad and most Y haplotypes are monophyletic, and it is also consistent with our previous work showing that recombination was suppressed in this region in the common ancestor with *P. wingei* ^22^.

**Figure 3.**
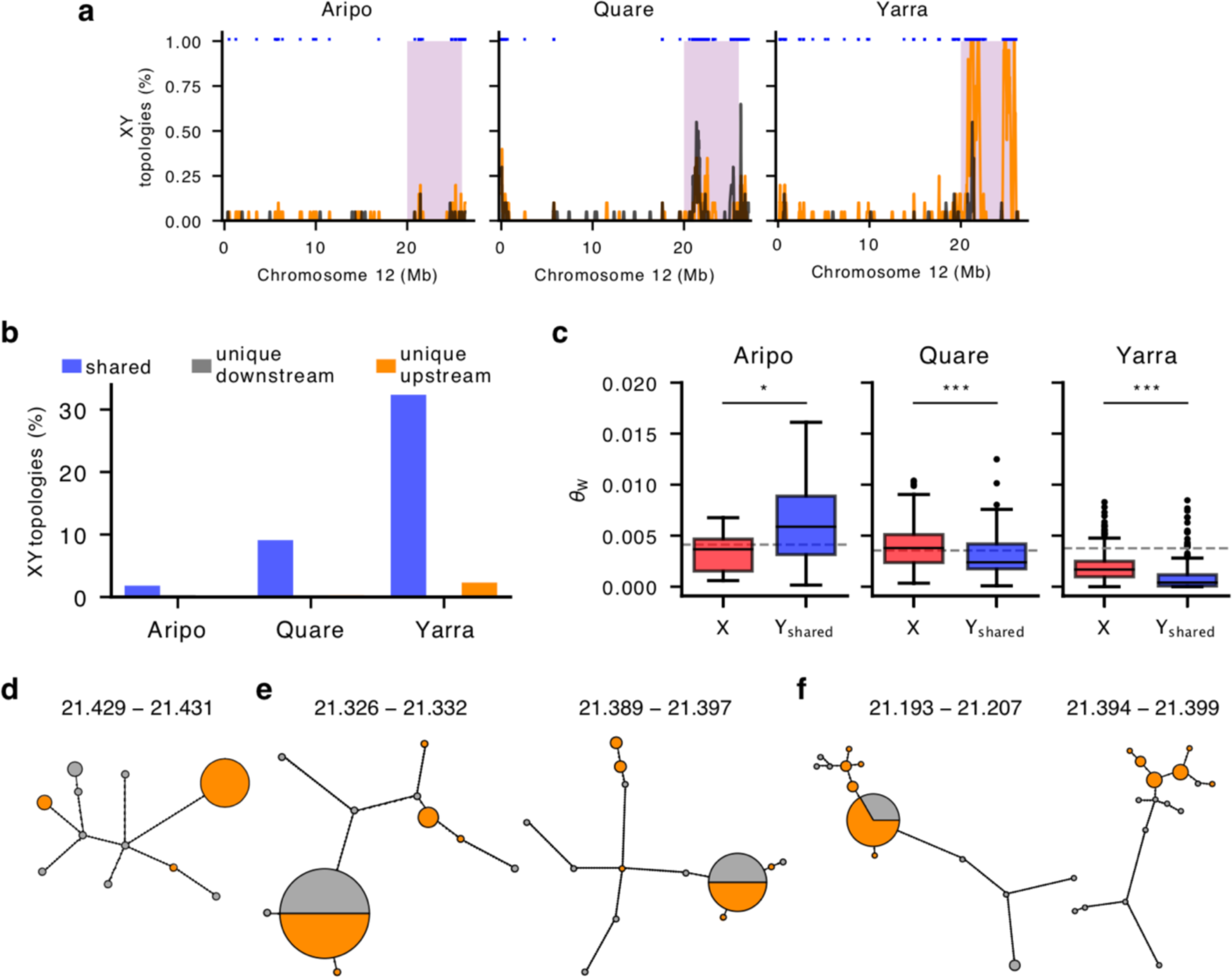
The guppy Y-chromosome shows exceptional genetic diversity. **a)** Distribution of tree topologies with separate clustering of X and Y SNPs (X-Y topology) on Chromosome 12 for downstream (black) and upstream (orange) populations in Aripo, Quare and Yarra watersheds. Stratum I is shaded in pink. Trees were inferred from phased alignments in non-overlapping windows of 100 SNPs with the Neighbour Joining method, see Materials and Methods for more details. The blue markers at the top indicate shared Y haplotypes between populations. **b)** Barplots showing the proportion of trees compatible with an XY sex chromosome system in Stratum I. Shared Y haplotypes were present in both populations and in > 66% of males in at least one population. **c)** Genetic diversity (Watterson’s theta estimator) of X and Y haplotypes within Stratum I in the Aripo, Quare and Yarra rivers. Y haplotypes were included only if found in both populations (Y_shared_) and were consistent with an XY topology in at least one of them. The dashed line indicates the estimated diversity for 1000 randomly sampled autosomal locations. ***p-value < 0.001, *p-value < 0.05. **d-f)** Networks of Y haplotypes in Aripo (d), Quare (e) and Yarra (f) rivers. Only networks for which the maximum number of Y haplotypes could be identified in both populations are shown. The circle area is proportional to haplotype frequency with the smallest circles representing single haplotypes, and branch lengths connecting haplotypes are proportional to the number of SNPs between haplotypes. Haplotypes from downstream populations in grey and upstream in orange. The approximate location (in Mb) of each region is indicated above the network.

We isolated shared male X- and Y-linked haplotypes from the gene trees described above and estimated nucleotide diversity for each aligned region. In comparison to the X chromosome, Y diversity was marginally higher in the Aripo watershed (θ_X_ = 0.003301, θ_Y_ = 0.006854, *P* = 0.010, Kruskal-Wallis test), and was significantly lower in Quare and Yarra (Quare θ_X_ = 0.003976, Quare θ_Y_ = 0.003143, Yarra θ_X_ = 0.001786, Yarra θ_Y_ = 0.001327, *P* < 0.001, Kruskal-Wallis test) (Figure 3c). Under a neutral model, diversity is expected to be proportional to the relative number of each chromosome in the population, therefore X diversity is expected to be 3/4 of autosomal diversity and Y diversity is expected to be 1/4 of that in the autosomes. Furthermore, linkage effects will deplete non-recombining regions of diversity, as Y chromosomes typically exhibit far less than 1/4 autosomal diversity ^21^. Both the X to autosome (X/A) as well as Y to autosome (Y/A) estimates depart from these neutral expectations. Y/A diversity was higher than expected in all three rivers, approaching ∼0.35 of that observed in the autosomes in Yarra and up to 1.66 in Aripo. In contrast, the X/A diversity was close to the expected 0.75 in Aripo (0.80), but was considerably higher in Quare (1.12) and considerably lower in Yarra (0.47), suggesting that selection and/or demography may be operating differentially in males and females.

To further explore the diversity and evolutionary history of the guppy Y chromosome, we built haplotype networks for the Y-linked regions for which Y haplotypes were shared across both populations (Figure 3d-f, Supplementary Figure S7; Supplementary Figure S8). We focused on the regions with haplotypes present in all or most males (Figure 3d-f) as these are the most informative. For both Quare and Yarra, this included two regions on Stratum I, between 21.3 Mb and 21.4 Mb, and one region in Aripo at ∼21.4 Mb. Except in Aripo where the most frequent haplotype is only present in upstream males, dominant haplotypes are shared between downstream and upstream populations. These dominant haplotypes include less than half of the male sequences sampled, and many of the Y haplotypes are observed in only one or two males, emphasizing the genetic diversity of Y chromosomes and the presence of low frequency Y-linked haplotypes in the guppy.

The number of mutational steps separating haplotypes of the same population was considerably higher in downstream populations with an average of 10.67, 17.14 and 14.5 mutations in Aripo, Quare and Yarra, respectively. In contrast, in the upstream populations there were 2.8 and 1.67 mutations for Quare and Yarra, respectively (the three upstream haplotypes observed in Aripo are not directly connected). These findings indicate higher haplotype diversity in the downstream populations, which is in line with the distribution of male-linked SNPs between the two populations. Some of the downstream Y haplotypes branch directly from the upstream population and may represent episodes of downstream migration.

### Characterisation of male-specific sequence

To identify the ancestral region of the Y chromosome in *P. reticulata*, we searched for 21 bp k-mer sequences unique to males and absent in all females (hereafter Y-mers) in each of our populations. We have previously used a Y-mer based approach to identify a small ancestral Y region across several *Poecilia* species ^22,23^. In order to evaluate the false positive rate (type-I error) for our sample sizes, we compared the number of Y-mers to female-specific k-mers (Supplementary Figure S9a). The latter can be used as a control because sequence unique to females is theoretically absent in a male heterogametic species. The false positive rate was <5% in all populations for Y-mers found in at least 6 males (Supplementary Figure S9b), so these thresholds were used for subsequent analyses.

Altogether, the number of population-specific Y-mers comprised ∼93% of the total number of Y-mers found in our dataset (Figure 4a), which suggests two non-mutually exclusive scenarios in the evolution of the Y chromosome in the guppy. The Y chromosomes have been evolving independently in each population since their split,expected if there were a large number of Y haplotypes at colonisation that vary in abundance across watersheds ^45^. Supporting this hypothesis, we also recovered only a very small number of Y-mers shared across the watersheds (13 Y-mers; Figure 4b), indicating a very dynamic nature of the Y chromosome. Alternatively, Y-specific sequence could also be accumulating separately within each population after recombination was suppressed in the ancestor. In line with this, most Y-mers map to repetitive rather than unique sequence (Supplementary Figure S10). This suggests that the build-up of repetitive sequence could be a major driver of sex chromosome divergence in the guppy, as it has been observed cytogenetically in *P. wingei* and to a less extent also in *P. reticulata* ^26,32^. Nonetheless, Stratum I was enriched for uniquely aligned Y-mers (Figure 4c), reinforcing again that this region is likely linked to the SDR and represents the region of ancestral recombination suppression.

**Figure 4.**
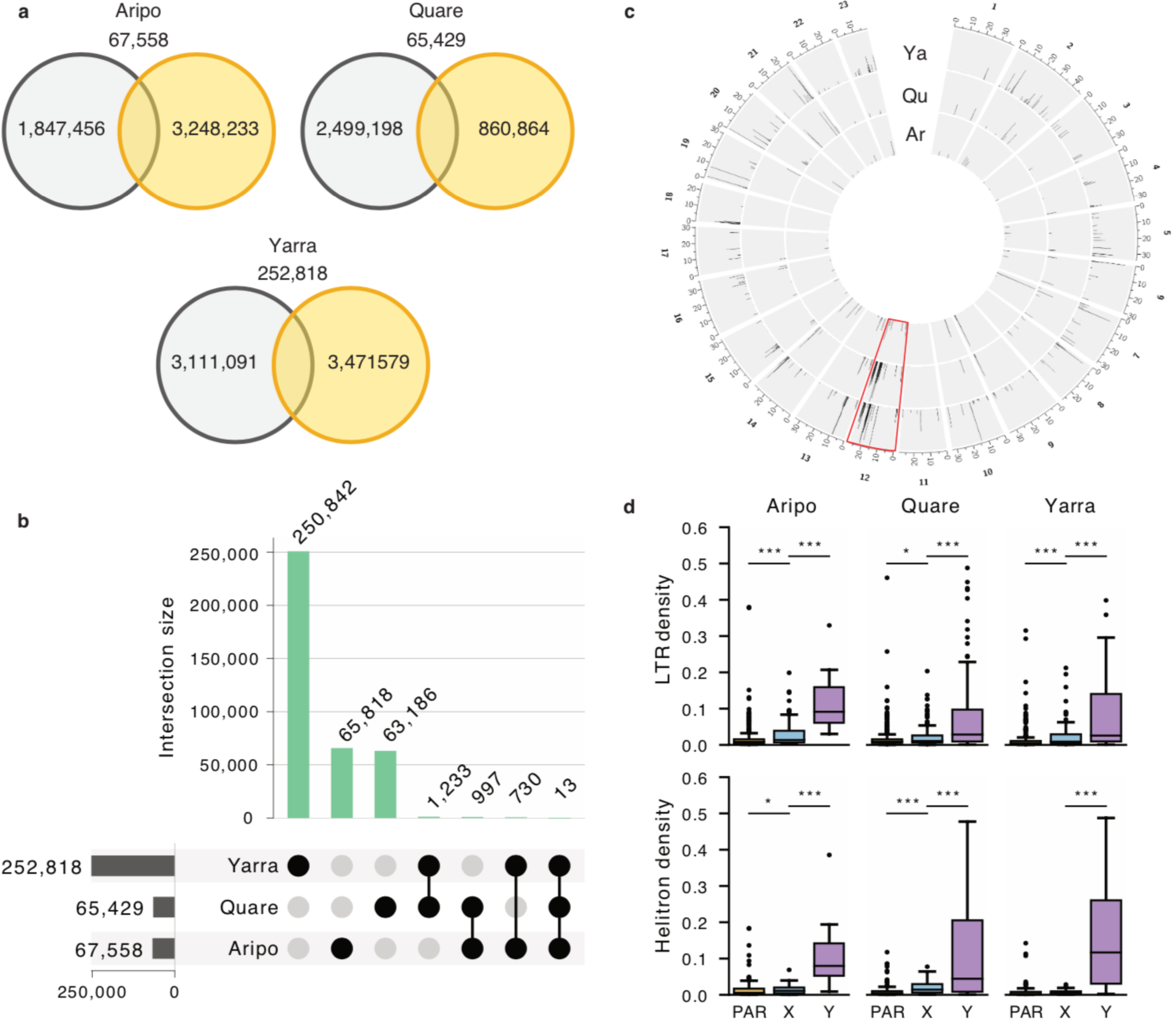
Characterisation of Y-linked sequence in the guppy. **a)** Venn diagrams showing the number and distribution of Y-mers among upstream (orange) and downstream (black) populations, along with overlapping Y-mers, **b)** Y-mers in the guppy are mostly river-specific, with only a limited number shared between watersheds. **c)** Circos plot showing the alignment of Y-mers to the reference genome of each population. Only Y-mers with unique alignments are shown. The sex chromosome (Chromosome 12) is highlighted in red. The region with the highest number of Y-mers aligned on Chromosome 12 overlaps with Stratum I (20 – 26 Mb). Ar: Aripo, Qu: Quare, Ya: Yarra. **d)** Transposable elements (TEs) have accumulated significantly more on the Y chromosome. Boxplots showing the density of LTRs and helitrons (total sequence of TEs in every non-overlapping 50 kb window) in scaffolds enriched with Y-mers.Statistics were calculated for the Y-linked scaffolds (Y), the region of the X chromosome homologous to the SDR (X) and to the pseudo-autosomal region (PAR). ***p-value < 0.001, **p-value < 0.01, *p-value < 0.05.

We extracted 10x Genomics Y haplotypes (megabubbles) enriched for Y-mers (see Materials & Methods) to further characterize Y-specific sequence. The density of transposable elements (TEs), measured as base-pairs of TE per 50 kb non-overlapping windows, was significantly higher in the candidate Y scaffolds, in comparison with both the PAR and the X-linked region of the sex chromosome (*P* < 0.001, Kruskal-Wallis test; Supplementary Figure S11a). In particular, TE enrichment was largely driven by Long Terminal Repeats (LTRs) and Helitrons (DNA transposons with a rolling-circle replication mechanism) that were markedly abundant in the Y scaffolds of all watersheds (Figure 4c, Supplementary Figure S11b). In addition, we also annotated a total of 193 potentially Y-linked genes, most of which are single copy (Supplementary Table S6). This estimate is, however, likely to include an unknown but probably considerable proportion of hitchhiking genes in these scaffolds. We could identify a homolog in the female guppy reference annotation for 69% of the genes, including in 19 out of the 22 genes common in at least two rivers. Six of these genes (htr1a-B, C6, C7, GHR, Nim1K and UNC13B) have been previously associated with the guppy SDR and mapped to a duplicated segment between Chromosome 9 and Chromosome 12 ^46^.

## Discussion

We used linked-read sequencing to assemble genomes for multiple individuals in downstream and upstream population pairs across three rivers in order to assess the divergence and diversity of the guppy Y chromosome. After replicating our previous identification of an ancestral non-recombining Y region shared across all watersheds, as well as greater overall sex chromosome divergence in replicate upstream compared to downstream populations ^15^, we identified Y-specific sequence in order to assess the degree of male haplotype diversity. We find that, contrary to the expected depletion of genetic variation that accompanies Y chromosome divergence due to the combined effects of linkage and purifying selection ^21^, the Y chromosome in guppies displays a remarkable level of diversity.

### Ancestral region of recombination suppression

By necessity, studying the initial stages of sex chromosome divergence requires studying sex chromosome systems with very low differentiation between the X and Y. This can be difficult given the subtlety of molecular signals that have barely begun to accumulate. It is perhaps not surprising then that our initial findings ^15,22^ have been challenged by others ^16,18^. In particular, Bergero et al. ^16^ did not observe any signal of Y degradation in Stratum I, nor increased Y divergence in upstream populations (Stratum II).

We have both replicated our previous results (Supplementary Figure S2, ^15,22^) using the same F_ST_ approach advocated by Bergero et al. (2019), and expanded it using several other lines of evidence in conjunction with our population-specific genome assemblies. We find consistent evidence of sex chromosome divergence within Stratum I, both due to reduced read mapping in males (Supplementary Figures S2 and S3) as well as the accumulation of male-linked SNPs (Figure 2), Y-mers, and repetitive elements (Figure 4). We observe sequence divergence particularly in two regions, 21-22 Mb and 25-26 Mb, where we see an excess of male-specific SNPs as well as an increase in gene tree topologies compatible with an XY sex chromosome system where Y chromosomes form a single cluster (Figure 3). These multiple concordant signatures of Y degeneration are common to all six wild populations analysed here, as well as recent comparative studies suggesting that the region of recombination suppression is ancestral to *P. reticulata* and its sister species, *P. wingei*, and therefore predates the colonisation of Trinidad ^22–24^.

We also again find evidence of Stratum II (Figure 1) ^15,22^, with greater male-female F_ST_ in all three replicate upstream populations compared to their downstream pair, consistent with convergence evolution of recombination suppression. After correcting for the structural difference between the reference genome and our populations (Supplementary Figure S1), elevated male-female F_ST_ in upstream populations extends over a greater length of Chromosome 12 than our previous estimates ^15^, and encompasses nearly the entirety of Chromosome 12 proximal to Stratum I.

### Y chromosome diversity

Within Stratum I, which achieved recombination suppression prior to the colonization of Trinidad ^23,24^, we observe a remarkable diversity of Y chromosome haplotypes, which is counter to the general expectations of strong sweep effects and low haplotype diversity expected of non-recombining Y regions. Instead, Y-specific sequence varies extensively across populations (Figure 4), and is supported by estimates of nucleotide variation (Figure 2, Figure 3c) as well as by haplotype networks (Figure 3d-f). Nucleotide diversity of Y haplotypes was also higher than expected from neutral models (Figure 3c). However, these models assume complete hemizygosity in males (for an XY system) which does not seem to be the case in the guppy Y chromosome (this study; Wright et al., 2017; Darolti et al., 2019, #86741; Bergero et al. 2019).

This diversity is initially puzzling, as non-recombining Y regions are typically depleted of variation via linkage effects and background selection. However, upon reflection our observation is consistent with organismal reports of Y haplotype diversity within the degenerated region. Specifically, YY males are viable only when Y chromosomes from different lineages are combined ^25,47^, suggesting both that many Y chromosome haplotypes contain recessive lethal variants and that the exact Y chromosome complement of these recessive lethal mutations varies across male lineages. Such high genetic diversity also matches well with the extraordinary phenotypic variation in male guppy colouration. Numerous reports in natural guppy populations suggest Y-linkage of many male colour traits and a variety of Y-linked colour combinations within populations ^13,19,48^, which together paint a picture of substantial Y diversity.

Recombination suppression is often thought to result once a large inversion fixes on the X or the Y chromosome ^4,49^. However, we do not observe an inversion (Supplemental Fig S5) in the non-recombining region of the Y chromosome, even though our use of long linked-read sequence from multiple individuals offers vastly increased power to detect chromosomal rearrangements relative to more traditional short-read sequencing. This lack of inversion could account for the marked differences between populations and rivers as well as the remarkable diversity that we observe in Y haplotypes. Because inversions are rare events, fixation of a Y inversion as a mechanism to achieve recombination suppression would lead to a strong bottleneck and a limited number of Y haplotypes. If recombination is curtailed through means other than an inversion, this bottleneck would not occur, leading to substantial initial haplotype diversity within the non-recombining region.

Several recent studies have indeed suggested that recombination suppression can proceed via means other than inversions ^50–53^ and also that inversions may follow recombination arrest in cases where they are not the cause of the initial recombination suppression ^5^. Our data are consistent with this. For example, we find a significant enrichment of male-linked SNPs in Stratum I, but relatively few are fixed in all downstream populations (Figure 2). This suggests that recombination suppression can be achieved via means other than an inversion, such as through differences in sex-specific recombination hotspots (heterochiasmy) ^54–56^ or from epigenetic variation ^57^. This, in turn, is expected to leave far greater Y chromosome diversity in the population.

However, despite the initial maintenance of Y haplotype diversity in the absence of an inversion, we would still expect the steady depletion of Y variation due to linkage effects and sweeps ^21^ without some other countering mechanism. Negative frequency dependent selection due to female preference for rare Y-linked male patterns ^58^ might act to maintain multiple male colour haplotypes on the Y chromosomes. Female guppies have long been known for their preference for rare or novel colouration phenotypes in males ^59,60^, and several experimental studies suggest that rare male coloration polymorphisms could be maintained by sexual selection via negative frequency-dependent selection ^58,61–63^. Our observation of an abundance of low-frequency Y haplotypes (Figure 3) is consistent with this hypothesis. Given that at least some genes controlling male colouration in the guppy are thought to be Y-linked ^13^, negative frequency-dependent selection could contribute to an increase in Y chromosome diversity and to the prevalence of multiple Y-linked genetic polymorphisms in the population. Future work linking specific colour combinations and Y haplotypes will be interesting to test the possible relationship between female preference and Y chromosome variation.

Finally, this and our previous work ^15^ supports a stronger association of sex-linked polymorphisms in upstream populations (Figure 1, Figure 2). Both the number of Y haplotypes unique to the upstream populations (not observed in downstream populations of the same watershed), and the genetic distance between upstream Y haplotypes were generally low (Figure 3b). This suggests that most of the upstream Y-linked variation is derived from the ancestral downstream populations, in agreement with other population genetic surveys ^64,65^.

## Conclusion

At the initial stages of sex chromosome divergence, population variation in the degree of sex-linkage is likely to maintain Y-linked alleles segregating in the male population ^66^, but the extent of such variation has been difficult to quantify in natural populations. Using phased X and Y haplotypes, our results show a remarkable population variation in the degree of sex-linkage in natural guppy populations, possibly due to the fact that recombination suppression is not based on an inversion. This, combined with other population-specific processes, such as frequency-dependent selection, act to maintain multiple Y polymorphisms segregating in natural guppy populations.

## Materials & Methods

### Sample collection

We collected wild *Poecilia reticulata* samples from three rivers (Yarra, Quare, Aripo) in the Northern Range Mountains of Trinidad in December 2016. A description of the habitats can be found in ^67^. From each river, we caught between 19 and 21 males and 20 females from both upstream and downstream populations. After dispatching samples, we immediately minced and placed heads in ethanol and flash froze tubes in liquid nitrogen. All samples were collected in accordance with national and institutional ethical guidelines.

### DNA extraction and sequencing

We prepared high molecular weight DNA samples from all males and females from each population. We extracted DNA from ∼25 mg of head tissue following an adapted 10x Genomics HMW DNA Extraction Protocol ^68^ and assessed molecular weight with the Agilent Femto Pulse system prior to library preparation. We then selected 10 males from each population with the highest molecular weights for 10x Genomics Chromium Sequencing. Because we were focused on phasing the X and Y chromosomes, linked reads were not required from female samples, however we still selected three females from each population with the highest molecular weights for 10x Chromium Sequencing. We used the 10X Genomics Chromium Genome Library Preparation Kit according to the manufacturer’s protocol (#CG00043 Chromium Genome Reagent Kit v2 User Guide), reducing the amount of starting DNA from the recommended 1.25 ng to 0.6 ng to account for the smaller genome size of *P. reticulata* (∼700 Mb) compared to the human genome, for which the protocol was optimized. The libraries were sequenced on an Illumina HiSeqX with 2 × 150 bp cycles using the v2.5 sequencing chemistry, resulting in an average of 346M reads per sample (range between 159M – 494M), representing an estimated average coverage of 68X (32X – 98X) (Supplementary Table S1).

For the remaining seven female samples in each population with lower molecular weight DNA, we generated additional Illumina sequencing data after extracting genomic DNA from ∼25 mg of head tissue using the DNeasy Blood & Tissue Kit (Qiagen) following the standard protocol. We prepared sequencing libraries from 1 ug DNA with the TruSeq PCRfree DNA Sample Preparation Kit, targeting an insert size of ∼350 bp according to the manufacturers’ instructions (guide #1000000039279). We sequenced resulting libraries on a Illumina HiSeqX with 2 × 150 bp cycles using the v2.5 sequencing chemistry, resulting in an average of 137M reads per sample 93M – 218M), representing 29X coverage on average (20X – 40X) (Supplementary Table S1).

### 10x Genomics de-novo assembly of genomes

To construct a reference genome specific for each river, we assembled linked reads de-novo for all our females with linked-read sequencing using Supernova v2.1.1 (10x Genomics). We then chose the best female assembly in each river based on scaffold N50 and phase block N50. Because Supernova can generate nearly identical haplotypes for the same sequence, we created a non-redundant assembly by removing smaller scaffolds with evidence of sequence overlap with longer scaffolds. For this, we aligned each assembly to itself with LAST v926 ^69^, using the NEAR DNA seeding scheme and post-masking of repeats (R11). To avoid false matches caused by repetitive sequences and paralogous scaffolds, we generated orthologous alignments with ‘last-split’, and discarded alignments with high proportions of masked sequence with ‘last-postmask’. We designated scaffolds as allelic variations in the assembly if they showed >90% sequence overlap and >95% sequence identity with other longer scaffolds.

We then used ARKS v1.0.2 ^70^ to increase the contiguity of each of our non-redundant assemblies. ARKS uses a k-mer approach to infer graph edges by determining the Chromium barcodes associated with the best-matching contig end for each read and selecting the contig end with the largest fraction of k-mer overlap. We ran ARKS with LINKS v1.8.6 ^71^ using default parameters and a Jaccard index = 0.5. Because different starting assemblies can have different optimal k-mers, we tested a range of k-mer values from 40 – 100 with increases of 20. As all k-mers tested produced a longer assembly N50 than the original, we chose the k-mer value that generated fewer scaffold miss-assemblies relative to the guppy reference genome ^37^, as determined by QUAST v5.0.2 ^72^.

Finally, we scaffolded each of our best ARKS assemblies, using the guppy reference genome (NCBI accession GCF_000633615.1) ^37^ as backbone, with RaGOO v1.02 ^73^. This process orders and orients assembled scaffolds relative to the chromosome-level reference genome. We ran RaGOO with a gap length of 100 and a minimum grouping confidence score = 0.3 instead of the default 0.2 in order to increase the precision of localised scaffolds. In general, only ∼1% (∼7 Mb) of the total genome size for each assembly could not be localised in the reference genome, which indicates a near complete placement of the assembly scaffolds.

In order to verify the scaffolding made by RaGOO, we performed pairwise synteny analyses between the assemblies and the NCBI guppy reference genome used in the scaffolding procedure. We made alignments with LAST v926 ^69^ using the NEAR DNA seeding scheme, post-masking repeats (R11) using a sensitive search with parameters ‘-m50 -C2’. We identified orthologous alignments with ‘last-split -m1’, discarding alignments comprised mostly of masked sequence with ‘last-postmask’. Although all of the three genomes showed strong co-linearity with the guppy NCBI reference genome, we identified an inverted segment on Chromosome 12 (between 0 – ∼9.9 Mb) that appears to have been translocated to ∼10.8 Mb. In order to verify that this rearrangement was not an artefact created during the genome assemblies, we independently aligned the 10x Genomics barcoded reads from each of the three female genomes, representing each river, to the guppy reference genome ^37^ using Long Ranger v2.2.2. From these read alignments, we identified clear breakpoints in all three female genomes for which the reads surrounding the breakpoints lack the expected pair orientation and/or are soft-clipped at the breakpoint positions (Supplementary Figure S1b,c). For consistency and direct comparison between the three rivers, we repeated the assembly scaffolding step above for the Quare and Yarra genomes using the Aripo genome as backbone, as the long scaffolds in the latter were correctly assembled for this inversion. This approach resulted in highly contiguous alignments without any evidence for an inversion in any of the female genomes.

### Pre-processing of sequencing reads

We used FastQC v0.11.5 ^74^ to assess read quality and used BBTools v38.34 “bbduk” ^75^ to remove adapter sequences, trim regions with average quality scores <Q10 and remove PhiX-174 spike-in control reads. After filtering, we excluded read-pairs from downstream analyses if either read had an average quality score <Q10 or was <35bp. In order to make both 10x Genomics and Illumina data sets comparable, we also pre-processed 10x Genomics reads as above after first trimming barcodes and performing barcode error correction with the Long Ranger v2.2.2 Basic pipeline.

### Read alignment and genotype calling

We aligned all pre-processed reads with BWA v0.7.15-r1140 using the MEM algorithm ^76^ and default options and processed the resulting SAM/BAM files with SAMtools v1.9 ^77^, flagging duplicated reads with biobambam v2.0.87 ^78^ after alignment.

We called genotypes within each river using Freebayes v1.3.1-16-g85d7bfc ^79^ with the following parameters: --min-repeat-entropy 1 --no-partial-observations --use-mapping-quality --min-mapping-quality 3 --min-base-quality 13 --skip-coverage 40000 --genotype-qualities --strict-vcf. Reads marked as duplicated and secondary alignments are excluded by default in Freebayes. We used ‘vcfallelicprimitives’ from vcflib v09df564 ^80^ to decompose complex variants into canonical SNP and indel representations and Vt v0.5772-60f436c3 ^81^ to normalize the resulting variants. We then used vcflib and BCFtools v1.9 ^82^ to perform a series of filters in order to exclude low quality variants and variants likely arising from copy number variations or paralogous sequences not present in the reference genomes ^83^. Specifically, we excluded variant sites with an alternate allele observation on the forward and reverse strands supported by <3 reads, variants with quality <30, heterozygous variants with depth <4 and homozygous variants with quality <50 and depth <4. If heterozygous, a variant was also excluded based on a maximum depth filter if the variant quality was <2x average depth and the maximum genotype depth was higher than the average depth + 3√(average depth) ^83^. Indels and single nucleotide polymorphisms (SNPs) within 3 bp of an indel were also excluded as the latter are difficult to ascertain with confidence. Finally, we filtered variants with an allele balance in the 10th percentile, but included fixed variants, and masked individual sample heterozygous genotypes if the corresponding genotype had <3 supporting reads for the reference and alternate allele. For all downstream analyses we only considered SNP variants with ≤10% missing genotypes.

### Degeneration and divergence analysis

Using our female genome assemblies, we expect that any regions of Y degeneration will result in female-biased read depth, as highly diverged Y reads will not map to the X chromosome, an approach previously implemented for guppies ^15,22^. We therefore assessed M:F read depth for each population separately by aligning pre-processed Illumina and 10x Genomics reads to the respective female river-specific genome assembly. We calculated per-site coverage with the SAMtools ‘depth’ command after applying stringent filtering criteria to exclude duplicated reads, reads with secondary alignments, non-unique alignments (with the XA or SA tags in the BAM file) and reads with a high number of mismatches to the reference genome (mapping quality <Q10). We restricted coverage calculations to the 23 assigned linkage groups in the guppy genome. We then calculated the effective coverage value as the median per site coverage in non-overlapping windows of 50 kb. To account for differences in the overall coverage between individuals, we normalized coverage data based on the median genomic coverage of each individual.

For regions of the X and Y chromosomes that have diverged, but still show little evidence of Y degeneration, we expect an increase in male compared to female diversity, as Y-linked reads will map to the female genome assembly, but will carry male-specific SNPs ^22,38^. We previously observed elevated male SNP density ^15^ in each of our upstream populations of *P. reticulata* across a large proportion of the sex chromosome compared to downstream populations. To investigate the extent of male to female divergence, we used Hudson’s method ^84^, as implemented in the Python library scikit-allel v1.2.1 ^85^, to calculate male:female F_ST_ for biallelic sites in non-overlapping windows of 50 kb. Windows of 10 kb produced qualitatively similar results (data not shown). Prior to F_ST_ calculation, we excluded singleton sites and sites with ≥20% of missing data in each sex.

### Phasing and analysis of Y haplotypes

We used the phasing information from linked-reads sequencing of 10 males and 3 females from each population to aid in phasing all our samples. 10x Genomics samples were assembled and phased using the 10x Genomics Long Ranger v.2.1.2 suite with the wgs option and using FreeBayes for variant calling. Briefly, reads that share the same barcode were sequenced from the same original long input DNA molecule, and this information is used to link the reads into large haplotype blocks. This is expected to generate a high quality phasing for each individual sample. To make the variation data between the 10x Genomics single-sample VCF and the pooled variants from each watershed as close as possible, we re-genotyped all individuals using the BAM files generated by Long Ranger. Genotyping was performed with FreeBayes and filtered for SNPs as detailed above.

The full phasing of all genotypes was then performed in two steps. First, we used WhatsHap v0.19.dev156+g1564a9f ^86^ for read-backed phasing. For the 10x Genomics samples, this simply maps the phase sets from the Long Ranger VCF to the river-specific genotyping VCF. In the case of the Illumina sequencing female samples, WhatsHap runs its full algorithm that uses the sequencing reads to reconstruct haplotypes. Secondly, we used SHAPEIT4 v4.1.2 ^87^ to computationally phase all samples. SHAPEIT4 was run to use the phase sets from WhatsHap with the recommended expected error rate of 0.01% and adjusting the default parameters for sequencing data (--use-PS 0.0001 --sequencing). To improve the phasing accuracy, we increased the number of iterations to ‘10b,1p,2b,1p,2b,1p,2b,1p,10m’ and also increased the number of conditioning neighbours to 8 (--pbwt-depth 8).

Y haplotypes can be identified phylogenetically using gene trees from the phased genotypes if they form a single monophyletic clade. For this, we divided the genome into non-overlapping windows of 100 SNPs and generated outputs in fasta format for the two inferred haplotypes of each individual with the BCFtools v1.9 consensus command. We used SNP windows for better resolution of regions with elevated SNP density. We then converted the fasta alignments at each window to the phylip format and built gene trees using FastME v 2.1.6.1 ^88^ with the parameters --method=BIONJ --dna=F84 --spr. For a given gene tree, we identified the Y chromosome haplotypes if a clade was composed exclusively of male individuals and included >66% males (all but three males) in each population (downstream and upstream populations). Because we are using river-specific references genomes, gene trees were computed separately in each river.

Nucleotide diversity was estimated using the Watterson’s theta estimator for the male Y haplotypes and the correspondingly alternative haplotypes (inferred as the X haplotypes). To obtain an autosomal diversity estimate we used a single arbitrarily chosen male haplotype from 1,000 randomly sampled autosomal gene trees (excluding Chromosome 12). To compute haplotype networks, we used the haploNet method with default parameters from the package pegas ^89^.

### Y-mer mining

k-mer refers to all the possible substrings of length k that are contained in a genome, and have been useful in identifying sex-specific (Y chromosome) sequence in a range of organisms ^90–92^, including guppies ^23^, by comparing male and female k-mer profiles. Male-specific k-mers, referred to here as Y-mers, most likely represent Y chromosome sequence, and Y-mers can be used to distinguish reads, read pairs, or scaffolds likely to be Y-linked. We used JELLYFISH v2.2.6 ^93^ to count the number of 21 bp canonical k-mers in the trimmed and filtered pre-processed reads for the male and female samples for each of our six wild guppy populations. We sorted and filtered k-mers, rejecting k-mers with observed counts lower than 3 in any individual, as these are likely sequencing errors. We then combined k-mer profiles for all samples of the same sex within each population, and identified male-specific k-mers absent from all female samples (Y-mers) with >5 average counts across males.

To identify scaffolds enriched for Y-specific sequence, we used Bowtie v1.2.3, without allowing for mismatches and reporting all alignments (-f -v 0 --all -l 21), to map Y-mers to the Supernova assemblies of each male genome. We used the megabubbles output in Supernova because in this output style Supernova generates an individual FASTA record for each homologous phased haplotype without mixing maternal and paternal alleles in the same sequence. Scaffolds with ≥100 Y-mers and more Y-mers than the homologous haplotype (if identified) were selected for further analysis. To remove possible redundancy in the resulting scaffolds, as the same region could be identified in scaffolds from different males, we clustered scaffolds within watersheds with a sequence identity threshold ≥90% and ≥50% of sequence overlap as calculated from BLASTN v2.5.0+ with parameters -dust yes -evalue 0.000001 -max_target_seqs 100000. The longest scaffold of each cluster with at least 5 samples, assuming that a Y-linked region could be absent from one sample due to insufficient coverage for assembly, was used for annotation.

### Annotation of Y-linked scaffolds

Candidate Y-linked scaffolds from the 3 watersheds were pooled together for annotation. Annotation was performed with MAKER v2.31.10 ^94^. We ran the MAKER pipeline twice: first based on a guppy-specific repeat library, protein sequence, EST and RNA sequence data (later used to train *ab-initio* software) and a second time combining evidence data from the first run and *ab-initio* predictions. We created a repeat library for these scaffolds using *de-novo* repeats identified by RepeatModeler v1.0.10 ^95^ which we then combined with Actinopterygii-specific repeats to use with RepeatMasker v4.0.7 ^96^. Annotated protein sequences were downloaded from Ensembl (release 95) ^97^ for 8 fish species: *Danio rerio* (GRCz11), *Gasterosteus aculeatus* (BROADS1), *Oryzias latipes* (ASM223467v1), *Poecilia latipinna* (1.0), *Poecilia mexicana* (1.0), *Poecilia reticulata* (1.0), *Takifugu rubripes* (FUGU5) and *Xiphophorus maculatus* (5.0). For EST, we used 10,664 tags from Dreyer et al. ^98^ isolated from guppy embryos and male testis. Furthermore, to support gene predictions we also used 2 publicly available libraries of RNA-seq data collected from guppy male testis and male embryos ^99^ and assembled with StringTie 1.3.3b ^100^. As basis for the construction of gene models, we combined *ab-initio* predictions from Augustus v3.2.3 ^101^, trained via BUSCO v3.0.2 ^102^, and SNAP v2006-07-28 ^103^. To train Augustus and SNAP, we first ran the MAKER pipeline a first time to create a profile using the protein and EST evidence along with RNA-seq data. Both Augustus and SNAP were then trained from this initial evidence-based annotation. Functional inference for genes and transcripts was performed using the translated CDS features of each coding transcript. Protein sequences were searched with BLAST in the Uniprot/Swissprot reference dataset in order to retrieve gene names and protein functions as well as in the InterProscan v5 database to retrieve additional annotations from different sources.

## Acknowledgements

This work was made possible by the European Research Council (grant agreement 680951) to J.E.M, who also gratefully acknowledges additional support from a Canada 150 Research Chair and the Natural Sciences and Engineering Research Council of Canada. We thank Y. Lin, L. Fong, W. van der Bijl, and D. Metzger for helpful comments.

