## Supplementary Tables and Figures for "Divergence and Remarkable Diversity of the Y Chromosome in Guppies"

**Supplementary Table S1.** Sequencing information for each sample.

| Sample | Sex | River | Population | Method | Raw data | Filtered data | | |
|---|---|---|---|---|---|---|---|---|
| | | | | | # reads | # reads | % reads | Coverage* |
| ArH-F1 | Female | Aripo | Downstream | Illumina | 177,001,862 | 176,505,596 | 99.72 | 28.97 |
| ArH-F9 | Female | Aripo | Downstream | Illumina | 123,686,724 | 123,461,396 | 99.82 | 16.83 |
| ArH-F10 | Female | Aripo | Downstream | Illumina | 139,802,356 | 139,241,328 | 99.60 | 22.89 |
| ArH-F13 | Female | Aripo | Downstream | Illumina | 105,446,352 | 105,237,508 | 99.80 | 17.27 |
| ArH-F14 | Female | Aripo | Downstream | Illumina | 163,351,722 | 162,980,354 | 99.77 | 22.89 |
| ArH-F23 | Female | Aripo | Downstream | Illumina | 191,928,134 | 191,596,890 | 99.83 | 31.57 |
| ArH-F25 | Female | Aripo | Downstream | Illumina | 109,682,202 | 109,408,568 | 99.75 | 17.94 |
| ARH-F2 | Female | Aripo | Downstream | 10x | 408,970,038 | 402,683,964 | 98.46 | 37.71 |
| ArH-F11 | Female | Aripo | Downstream | 10x | 381,405,262 | 368,721,272 | 96.67 | 32.21 |
| ARH-F24 | Female | Aripo | Downstream | 10x | 426,713,902 | 420,185,200 | 98.47 | 38.33 |
| ARH-M1 | Male | Aripo | Downstream | 10x | 439,226,730 | 432,609,612 | 98.49 | 39.32 |
| ArH-M2 | Male | Aripo | Downstream | 10x | 252,352,686 | 246,637,432 | 97.74 | 26.50 |
| ArH-M5 | Male | Aripo | Downstream | 10x | 260,888,232 | 256,042,222 | 98.14 | 23.71 |
| ArH-M7 | Male | Aripo | Downstream | 10x | 355,642,886 | 348,246,874 | 97.92 | 35.55 |
| ArH-M9 | Male | Aripo | Downstream | 10x | 387,869,768 | 380,303,506 | 98.05 | 40.35 |
| ArH-M12 | Male | Aripo | Downstream | 10x | 288,489,532 | 283,211,948 | 98.17 | 26.85 |
| ARH-M16 | Male | Aripo | Downstream | 10x | 466,170,214 | 459,951,168 | 98.67 | 45.05 |
| ArH-M19 | Male | Aripo | Downstream | 10x | 485,992,400 | 471,350,870 | 96.99 | 35.21 |
| ArH-M21 | Male | Aripo | Downstream | 10x | 241,425,146 | 235,678,560 | 97.62 | 23.68 |
| ArH-M24 | Male | Aripo | Downstream | 10x | 302,563,204 | 295,221,194 | 97.57 | 26.84 |
| ArL-F5 | Female | Aripo | Upstream | Illumina | 125,681,718 | 125,115,518 | 99.55 | 21.00 |
| ArL-F8 | Female | Aripo | Upstream | Illumina | 145,627,362 | 145,164,072 | 99.68 | 24.14 |
| ArL-F9 | Female | Aripo | Upstream | Illumina | 146,020,928 | 145,499,770 | 99.64 | 24.39 |
| ArL-F15 | Female | Aripo | Upstream | Illumina | 139,996,196 | 139,510,334 | 99.65 | 23.47 |

| | | | | | | | | |
|---|---|---|---|---|---|---|---|---|
| ArL-F18 | Female | Aripo | Upstream | Illumina | 149,839,898 | 149,483,814 | 99.76 | 24.85 |
| ArL-F19 | Female | Aripo | Upstream | Illumina | 129,400,664 | 128,984,726 | 99.68 | 21.64 |
| ArL-F20 | Female | Aripo | Upstream | Illumina | 121,795,694 | 121,199,836 | 99.51 | 20.19 |
| ArL-F10 | Female | Aripo | Upstream | 10x | 331,020,806 | 326,156,754 | 98.53 | 30.38 |
| ArL-F16 | Female | Aripo | Upstream | 10x | 337,717,026 | 332,234,258 | 98.38 | 29.81 |
| ArL-F22 | Female | Aripo | Upstream | 10x | 281,639,728 | 275,613,302 | 97.86 | 26.56 |
| ArL-M1 | Male | Aripo | Upstream | 10x | 395,101,320 | 385,055,732 | 97.46 | 34.29 |
| ArL-M6 | Male | Aripo | Upstream | 10x | 344,463,254 | 336,457,206 | 97.68 | 29.02 |
| ArL-M11 | Male | Aripo | Upstream | 10x | 374,115,442 | 364,106,466 | 97.32 | 31.81 |
| ArL-M12 | Male | Aripo | Upstream | 10x | 307,371,902 | 296,634,782 | 96.51 | 27.48 |
| ARL-M15 | Male | Aripo | Upstream | 10x | 385,877,220 | 380,935,470 | 98.72 | 37.81 |
| ArL-M16 | Male | Aripo | Upstream | 10x | 369,068,090 | 360,681,320 | 97.73 | 35.27 |
| ARL-M17 | Male | Aripo | Upstream | 10x | 395,209,344 | 389,830,668 | 98.64 | 37.37 |
| ArL-M19 | Male | Aripo | Upstream | 10x | 354,769,736 | 346,126,668 | 97.56 | 28.64 |
| ArL-M24 | Male | Aripo | Upstream | 10x | 316,277,460 | 309,367,708 | 97.82 | 32.45 |
| ARL-M25 | Male | Aripo | Upstream | 10x | 348,601,062 | 343,193,820 | 98.45 | 32.86 |
| QuH-F4 | Female | Quare | Downstream | Illumina | 179,052,012 | 178,453,836 | 99.67 | 28.62 |
| QuH-F6 | Female | Quare | Downstream | Illumina | 106,006,310 | 105,684,712 | 99.70 | 17.02 |
| QuH-F7 | Female | Quare | Downstream | Illumina | 146,614,088 | 146,074,580 | 99.63 | 25.19 |
| QuH-F9 | Female | Quare | Downstream | Illumina | 125,867,070 | 125,295,106 | 99.55 | 21.45 |
| QuH-F12 | Female | Quare | Downstream | Illumina | 147,284,914 | 146,610,000 | 99.54 | 24.65 |
| QuH-F13 | Female | Quare | Downstream | Illumina | 147,789,606 | 147,211,156 | 99.61 | 25.05 |
| QuH-F22 | Female | Quare | Downstream | Illumina | 122,308,096 | 122,024,688 | 99.77 | 17.76 |
| QuH-F3 | Female | Quare | Downstream | 10x | 383,364,788 | 376,444,214 | 98.19 | 34.92 |
| QuH-F21 | Female | Quare | Downstream | 10x | 372,586,380 | 364,173,432 | 97.74 | 34.35 |
| QuH-F25 | Female | Quare | Downstream | 10x | 262,839,572 | 258,146,910 | 98.21 | 24.71 |
| QuH-M2 | Male | Quare | Downstream | 10x | 332,995,698 | 325,603,698 | 97.78 | 28.54 |
| QuH-M3 | Male | Quare | Downstream | 10x | 304,584,270 | 299,436,134 | 98.31 | 29.42 |

| | | | | | | | | |
|---|---|---|---|---|---|---|---|---|
| QUH-M4 | Male | Quare | Downstream | 10x | 397,440,770 | 391,344,378 | 98.47 | 34.47 |
| QUH-M9 | Male | Quare | Downstream | 10x | 397,506,462 | 391,924,012 | 98.60 | 40.17 |
| QuH-M10 | Male | Quare | Downstream | 10x | 306,774,330 | 300,809,780 | 98.06 | 29.66 |
| QuH-M13 | Male | Quare | Downstream | 10x | 159,301,776 | 156,661,362 | 98.34 | 17.66 |
| QuH-M17 | Male | Quare | Downstream | 10x | 334,204,944 | 328,118,800 | 98.18 | 32.04 |
| QuH-M19 | Male | Quare | Downstream | 10x | 329,744,366 | 324,159,304 | 98.31 | 31.40 |
| QuH-M22 | Male | Quare | Downstream | 10x | 339,581,552 | 332,344,120 | 97.87 | 28.45 |
| QuL-F4 | Female | Quare | Upstream | Illumina | 113,172,552 | 112,900,498 | 99.76 | 19.19 |
| QuL-F6 | Female | Quare | Upstream | Illumina | 123,311,704 | 122,950,174 | 99.71 | 20.49 |
| QuL-F7 | Female | Quare | Upstream | Illumina | 116,293,246 | 116,001,858 | 99.75 | 18.74 |
| QuL-F14 | Female | Quare | Upstream | Illumina | 218,161,778 | 217,313,576 | 99.61 | 35.62 |
| QuL-F15 | Female | Quare | Upstream | Illumina | 130,978,920 | 130,701,716 | 99.79 | 21.02 |
| QuL-F19 | Female | Quare | Upstream | Illumina | 92,666,384 | 92,425,842 | 99.74 | 15.16 |
| QuL-F23 | Female | Quare | Upstream | Illumina | 118,582,976 | 118,221,148 | 99.69 | 18.98 |
| QUL-F5 | Female | Quare | Upstream | 10x | 359,994,034 | 355,396,326 | 98.72 | 34.67 |
| QuL-F11 | Female | Quare | Upstream | 10x | 357,689,770 | 349,210,524 | 97.63 | 29.03 |
| QUL-F16 | Female | Quare | Upstream | 10x | 478,732,114 | 471,211,580 | 98.43 | 44.48 |
| QuL-M2 | Male | Quare | Upstream | 10x | 308,996,474 | 303,509,088 | 98.22 | 36.16 |
| QUL-M4 | Male | Quare | Upstream | 10x | 378,405,724 | 371,842,898 | 98.27 | 35.87 |
| QuL-M5 | Male | Quare | Upstream | 10x | 376,139,404 | 365,675,072 | 97.22 | 28.20 |
| QuL-M6 | Male | Quare | Upstream | 10x | 339,915,846 | 333,810,516 | 98.20 | 28.95 |
| QuL-M8 | Male | Quare | Upstream | 10x | 370,727,306 | 364,327,302 | 98.27 | 33.78 |
| QuL-M11 | Male | Quare | Upstream | 10x | 323,424,146 | 316,698,712 | 97.92 | 26.86 |
| QuL-M15 | Male | Quare | Upstream | 10x | 317,275,484 | 312,358,620 | 98.45 | 28.26 |
| QuL-M17 | Male | Quare | Upstream | 10x | 346,548,086 | 341,135,520 | 98.44 | 30.95 |
| QuL-M20 | Male | Quare | Upstream | 10x | 324,939,344 | 318,958,212 | 98.16 | 38.31 |
| QuL-M25 | Male | Quare | Upstream | 10x | 338,420,592 | 333,069,042 | 98.42 | 34.08 |
| YaH-F9 | Female | Yarra | Downstream | Illumina | 154,552,664 | 154,130,620 | 99.73 | 21.32 |

| | | | | | | | | |
|---|---|---|---|---|---|---|---|---|
| YaH-F13 | Female | Yarra | Downstream | Illumina | 137,843,852 | 137,496,702 | 99.75 | 20.34 |
| YaH-F14 | Female | Yarra | Downstream | Illumina | 142,201,016 | 141,656,306 | 99.62 | 19.50 |
| YaH-F15 | Female | Yarra | Downstream | Illumina | 152,591,576 | 152,311,356 | 99.82 | 22.21 |
| YaH-F18 | Female | Yarra | Downstream | Illumina | 130,309,702 | 129,898,602 | 99.68 | 19.13 |
| YaH-F21 | Female | Yarra | Downstream | Illumina | 133,816,960 | 133,394,266 | 99.68 | 18.42 |
| YaH-F24 | Female | Yarra | Downstream | Illumina | 122,636,722 | 122,188,898 | 99.63 | 16.68 |
| YaH-F1 | Female | Yarra | Downstream | 10x | 222,780,570 | 219,055,424 | 98.33 | 26.18 |
| YAH-F12 | Female | Yarra | Downstream | 10x | 356,192,978 | 350,570,380 | 98.42 | 34.30 |
| YaH-F19 | Female | Yarra | Downstream | 10x | 348,295,768 | 340,287,472 | 97.70 | 30.88 |
| YaH-M2 | Male | Yarra | Downstream | 10x | 236,620,736 | 231,584,160 | 97.87 | 23.60 |
| YaH-M5 | Male | Yarra | Downstream | 10x | 405,433,092 | 392,515,154 | 96.81 | 27.71 |
| YaH-M6 | Male | Yarra | Downstream | 10x | 298,369,238 | 282,064,166 | 94.54 | 19.68 |
| YaH-M10 | Male | Yarra | Downstream | 10x | 323,420,634 | 316,618,056 | 97.90 | 33.81 |
| YaH-M11 | Male | Yarra | Downstream | 10x | 295,137,132 | 286,247,122 | 96.99 | 21.57 |
| YaH-M16 | Male | Yarra | Downstream | 10x | 358,627,758 | 345,447,188 | 96.32 | 22.49 |
| YaH-M17 | Male | Yarra | Downstream | 10x | 328,438,330 | 316,504,572 | 96.37 | 23.90 |
| YaH-M18 | Male | Yarra | Downstream | 10x | 220,144,952 | 216,739,042 | 98.45 | 26.14 |
| YaH-M21 | Male | Yarra | Downstream | 10x | 326,897,482 | 315,901,204 | 96.64 | 23.42 |
| YaH-M23 | Male | Yarra | Downstream | 10x | 331,356,786 | 317,506,648 | 95.82 | 26.82 |
| YaL-F4 | Female | Yarra | Upstream | Illumina | 138,514,074 | 138,110,048 | 99.71 | 20.81 |
| YaL-F6 | Female | Yarra | Upstream | Illumina | 134,171,018 | 133,760,786 | 99.69 | 20.05 |
| YaL-F10 | Female | Yarra | Upstream | Illumina | 117,695,078 | 117,470,256 | 99.81 | 20.12 |
| YaL-F12 | Female | Yarra | Upstream | Illumina | 140,113,924 | 139,528,912 | 99.58 | 23.87 |
| YaL-F19 | Female | Yarra | Upstream | Illumina | 139,747,920 | 139,451,146 | 99.79 | 20.42 |
| YaL-F22 | Female | Yarra | Upstream | Illumina | 125,894,888 | 125,487,600 | 99.68 | 21.17 |
| YaL-F24 | Female | Yarra | Upstream | Illumina | 125,153,320 | 124,768,922 | 99.69 | 17.14 |
| YaL-F11 | Female | Yarra | Upstream | 10x | 391,722,230 | 382,605,508 | 97.67 | 35.15 |
| YaL-F13 | Female | Yarra | Upstream | 10x | 307,344,588 | 300,997,862 | 97.93 | 28.00 |

| | | | | | | | | |
|---|---|---|---|---|---|---|---|---|
| YAL-F20 | Female | Yarra | Upstream | 10x | 391,780,204 | 384,760,566 | 98.21 | 32.96 |
| YaL-M7 | Male | Yarra | Upstream | 10x | 363,941,054 | 357,078,016 | 98.11 | 32.47 |
| YAL-M8 | Male | Yarra | Upstream | 10x | 451,668,232 | 445,186,766 | 98.56 | 39.19 |
| YAL-M10 | Male | Yarra | Upstream | 10x | 494,223,386 | 483,699,440 | 97.87 | 40.37 |
| YaL-M11 | Male | Yarra | Upstream | 10x | 259,374,704 | 255,007,362 | 98.32 | 25.58 |
| YaL-M14 | Male | Yarra | Upstream | 10x | 409,708,944 | 398,129,722 | 97.17 | 29.28 |
| YaL-M16 | Male | Yarra | Upstream | 10x | 292,372,948 | 285,855,314 | 97.77 | 26.45 |
| YaL-M17 | Male | Yarra | Upstream | 10x | 267,080,378 | 262,237,792 | 98.19 | 26.12 |
| YaL-M18 | Male | Yarra | Upstream | 10x | 399,152,812 | 388,825,948 | 97.41 | 34.47 |
| YaL-M21 | Male | Yarra | Upstream | 10x | 368,029,652 | 360,600,418 | 97.98 | 34.95 |
| YaL-M23 | Male | Yarra | Upstream | 10x | 420,027,608 | 409,077,362 | 97.39 | 29.61 |
| YaL-M25 | Male | Yarra | Upstream | 10x | 332,547,264 | 325,775,770 | 97.96 | 29.89 |

* effective coverage of reference-guided alignments to river-specific reference genomes after read trimming and duplicate removal.

**Supplementary Table S2.** Assembly statistics for all 10x Genomics samples using Supernova *de-novo* assembler.

| Sample | Sex | Source | Valid barcode % | Effective coverage | Molecule length | Assembly size | Contig N50 | Scaffold N50 | Scaffolds 10kb+ | Scaffolds 1kb+ | Phase N50 |
|---|---|---|---|---|---|---|---|---|---|---|---|
| ArH-F2 | Female | Ar-D | 92.89 | 34.34 | 12,479 | 603,896,307 | 30,435 | 61,210 | 13,707 | 51,535 | 109,106 |
| ArH-F11 | Female | Ar-D | 94.61 | 29.87 | 11,873 | 570,483,834 | 25,621 | 49,550 | 16,200 | 56,809 | 67,120 |
| ArH-F24 | Female | Ar-D | 94.18 | 35.15 | 29,363 | 729,242,799 | 50,348 | 1,584,760 | 3,703 | 22,339 | 593,022 |
| ArH-M1 | Male | Ar-D | 93.99 | 35.56 | 22,003 | 672,922,408 | 48,757 | 479,207 | 4,774 | 22,456 | 369,523 |
| ArH-M2 | Male | Ar-D | 93.89 | 24.74 | 23,357 | 646,434,489 | 36,408 | 116,760 | 10,373 | 35,215 | 173,504 |
| ArH-M5 | Male | Ar-D | 94.15 | 21.48 | 20,319 | 648,310,764 | 39,976 | 295,457 | 6,592 | 29,422 | 266,082 |
| ArH-M7 | Male | Ar-D | 93.07 | 32.92 | 23,826 | 666,756,002 | 43,787 | 243,248 | 7,181 | 24,657 | 266,503 |
| ArH-M9 | Male | Ar-D | 93.59 | 37.82 | 24,794 | 677,369,114 | 51,770 | 638,371 | 4,382 | 24,866 | 500,136 |
| ArH-M12 | Male | Ar-D | 94.13 | 24.41 | 21,144 | 671,597,633 | 47,579 | 639,034 | 4,288 | 19,471 | 326,840 |
| ArH-M16 | Male | Ar-D | 93.63 | 41.17 | 22,003 | 677,689,639 | 51,850 | 1,280,894 | 3,288 | 20,004 | 802,655 |
| ArH-M19 | Male | Ar-D | 93.07 | 31.66 | nd | 84,351,704 | 15,431 | 16,164 | 5,101 | 132,798 | 2,874 |
| ArH-M21 | Male | Ar-D | 93.32 | 20.96 | 16,325 | 597,049,153 | 26,596 | 48,636 | 16,778 | 54,789 | 67,539 |
| ArH-M24 | Male | Ar-D | 94.35 | 24.94 | 9,926 | 600,957,212 | 28,856 | 45,523 | 17,703 | 49,508 | 56,221 |
| ArL-F10 | Female | Ar-U | 93.86 | 28.00 | 22,003 | 682,875,263 | 54,076 | 1,518,110 | 3,022 | 14,584 | 557,061 |
| ArL-F16 | Female | Ar-U | 93.78 | 26.84 | 33,418 | 683,530,463 | 54,203 | 3,333,357 | 2,517 | 13,828 | 1,079,421 |
| ArL-F22 | Female | Ar-U | 94.26 | 23.90 | 20,118 | 667,305,062 | 42,648 | 345,876 | 5,857 | 22,437 | 267,979 |
| ArL-M1 | Male | Ar-U | 94.25 | 31.45 | 18,032 | 670,441,308 | 40,452 | 173,277 | 8,260 | 25,177 | 176,037 |
| ArL-M6 | Male | Ar-U | 94.01 | 26.32 | 23,357 | 675,835,125 | 43,709 | 357,112 | 5,849 | 20,424 | 263,802 |
| ArL-M11 | Male | Ar-U | 94.44 | 29.05 | 24,794 | 673,017,375 | 42,428 | 229,772 | 6,736 | 22,192 | 206,621 |
| ArL-M12 | Male | Ar-U | 94.67 | 25.33 | 11,074 | 621,217,985 | 28,539 | 47,766 | 17,259 | 41,614 | 55,199 |
| ArL-M15 | Male | Ar-U | 93.09 | 33.94 | 26,848 | 685,292,827 | 52,226 | 2,842,056 | 2,832 | 16,074 | 887,754 |
| ArL-M16 | Male | Ar-U | 94.42 | 32.45 | 21,356 | 686,717,238 | 50,777 | 883,946 | 3,690 | 15,732 | 541,151 |
| ArL-M17 | Male | Ar-U | 92.92 | 33.62 | 14,061 | 670,913,526 | 44,741 | 211,057 | 6,905 | 22,347 | 162,071 |
| ArL-M19 | Male | Ar-U | 93.91 | 25.35 | 13,512 | 616,636,301 | 29,873 | 54,820 | 15,075 | 40,051 | 64,261 |

| | | | | | | | | | | | |
|---|---|---|---|---|---|---|---|---|---|---|---|
| ArL-M24 | Male | Ar-U | 93.75 | 30.23 | 6,869 | 580,898,802 | 29,105 | 42,628 | 17,901 | 49,183 | 55,408 |
| ArL-M25 | Male | Ar-U | 94.05 | 30.24 | 20,319 | 676,980,460 | 51,386 | 725,226 | 3,906 | 17,368 | 415,239 |
| QuH-F3 | Female | Qu-D | 94.02 | 32.20 | 12,233 | 615,355,903 | 30,730 | 59,296 | 14,154 | 50,604 | 105,695 |
| QuH-F21 | Female | Qu-D | 94.38 | 32.08 | 9,166 | 574,798,522 | 28,246 | 43,776 | 17,691 | 59,624 | 59,614 |
| QuH-F25 | Female | Qu-D | 94.13 | 24.74 | 28,500 | 641,956,330 | 39,791 | 319,579 | 7,006 | 28,515 | 307,182 |
| QuH-M2 | Male | Qu-D | 93.95 | 25.98 | 5,858 | 295,245,989 | 15,868 | 19,569 | 15,930 | 112,140 | 25,216 |
| QuH-M3 | Male | Qu-D | 93.55 | 26.69 | 11,639 | 559,237,759 | 28,649 | 48,359 | 16,102 | 61,755 | 66,441 |
| QuH-M4 | Male | Qu-D | 93.47 | 30.45 | 23,826 | 669,172,328 | 50,682 | 778,580 | 3,698 | 19,469 | 454,672 |
| QuH-M9 | Male | Qu-D | 93.56 | 36.89 | 18,032 | 673,996,478 | 53,012 | 791,350 | 3,736 | 19,521 | 432,210 |
| QuH-M10 | Male | Qu-D | 94.10 | 27.49 | 11,755 | 558,402,613 | 26,999 | 49,136 | 15,908 | 64,502 | 77,392 |
| QuH-M13 | Male | Qu-D | 94.32 | 16.29 | 24,064 | 577,989,203 | 26,391 | 137,986 | 9,804 | 48,823 | 161,594 |
| QuH-M17 | Male | Qu-D | 93.81 | 29.38 | 16,325 | 645,193,812 | 37,249 | 101,325 | 10,804 | 38,131 | 152,354 |
| QuH-M19 | Male | Qu-D | 93.71 | 28.59 | 11,410 | 552,950,240 | 27,686 | 48,242 | 16,282 | 64,242 | 74,587 |
| QuH-M22 | Male | Qu-D | 93.62 | 25.76 | 7,438 | 546,134,524 | 25,575 | 36,669 | 18,916 | 61,646 | 52,960 |
| QuL-F5 | Female | Qu-U | 92.81 | 30.86 | 35,474 | 685,901,629 | 53,492 | 5,195,008 | 1,935 | 9,980 | 752,633 |
| QuL-F11 | Female | Qu-U | 93.98 | 25.90 | 5,251 | 453,708,191 | 19,440 | 23,168 | 21,506 | 73,073 | 13,022 |
| QuL-F16 | Female | Qu-U | 94.13 | 40.02 | 16,163 | 682,908,655 | 53,788 | 732,948 | 3,061 | 11,812 | 301,380 |
| QuL-M2 | Male | Qu-U | 93.63 | 34.01 | 26,058 | 689,830,587 | 61,318 | 2,298,331 | 2,071 | 9,801 | 510,678 |
| QuL-M4 | Male | Qu-U | 94.05 | 32.64 | 26,848 | 681,774,619 | 55,402 | 2,186,889 | 2,101 | 11,233 | 563,437 |
| QuL-M5 | Male | Qu-U | 94.08 | 25.24 | 4,390 | 355,020,979 | 17,467 | 18,886 | 19,401 | 87,904 | 4,001 |
| QuL-M6 | Male | Qu-U | 93.98 | 26.16 | 25,545 | 674,390,728 | 48,692 | 579,889 | 3,951 | 13,599 | 300,025 |
| QuL-M8 | Male | Qu-U | 94.27 | 31.43 | 21,785 | 685,637,933 | 56,036 | 922,322 | 2,921 | 11,240 | 327,493 |
| QuL-M11 | Male | Qu-U | 94.17 | 24.32 | 11,755 | 632,789,117 | 32,662 | 54,421 | 15,250 | 34,413 | 60,085 |
| QuL-M15 | Male | Qu-U | 93.81 | 25.39 | 25,800 | 680,690,864 | 53,789 | 1,703,422 | 2,525 | 11,868 | 406,911 |
| QuL-M17 | Male | Qu-U | 94.01 | 27.94 | 26,582 | 687,822,904 | 56,629 | 2,561,222 | 2,242 | 11,099 | 524,879 |
| QuL-M20 | Male | Qu-U | 93.86 | 35.85 | 38,797 | 692,284,958 | 58,647 | 4,694,176 | 1,941 | 10,097 | 806,503 |
| QuL-M25 | Male | Qu-U | 93.82 | 31.27 | 36,187 | 688,491,763 | 59,049 | 4,135,504 | 1,999 | 10,226 | 760,282 |
| YaH-F1 | Female | Ya-D | 93.77 | 24.84 | 26,058 | 662,631,832 | 48,239 | 1,215,823 | 3,303 | 18,049 | 364,287 |

| | | | | | | | | | | | |
|---|---|---|---|---|---|---|---|---|---|---|---|
| YaH-F12 | Female | Ya-D | 93.42 | 32.61 | 27,387 | 670,254,389 | 45,107 | 549,157 | 5,056 | 20,604 | 306,641 |
| YaH-F19 | Female | Ya-D | 94.30 | 28.78 | 4,613 | 216,742,755 | 14,626 | 16,251 | 13,267 | 116,625 | 4,554 |
| YaH-M2 | Male | Ya-D | 94.47 | 21.87 | 7,438 | 514,720,685 | 25,053 | 36,636 | 18,026 | 64,522 | 46,584 |
| YaH-M5 | Male | Ya-D | 94.39 | 24.49 | 13,922 | 661,982,631 | 33,347 | 78,737 | 12,837 | 36,672 | 116,081 |
| YaH-M6 | Male | Ya-D | 94.58 | 17.94 | nd | 106,720,199 | 14,669 | 14,920 | 6,988 | 144,481 | 8,184 |
| YaH-M10 | Male | Ya-D | 93.44 | 31.53 | 14,778 | 638,343,761 | 34,690 | 84,743 | 11,527 | 38,998 | 118,361 |
| YaH-M11 | Male | Ya-D | 94.39 | 18.32 | 9,730 | 635,051,176 | 30,379 | 63,598 | 13,542 | 35,381 | 72,759 |
| YaH-M16 | Male | Ya-D | 94.62 | 19.53 | 6,343 | 335,196,646 | 17,259 | 22,059 | 16,553 | 95,458 | 32,997 |
| YaH-M17 | Male | Ya-D | 94.49 | 21.71 | 7,438 | 594,225,669 | 27,052 | 44,255 | 17,478 | 49,231 | 54,745 |
| YaH-M18 | Male | Ya-D | 93.47 | 24.54 | 33,087 | 655,984,201 | 48,911 | 3,381,888 | 2,522 | 17,986 | 652,943 |
| YaH-M21 | Male | Ya-D | 94.52 | 20.70 | 9,351 | 605,902,349 | 27,637 | 48,967 | 16,418 | 48,044 | 58,163 |
| YaH-M23 | Male | Ya-D | 93.57 | 24.71 | nd | 133,227,602 | 15,422 | 15,974 | 8,145 | 131,841 | 3,866 |
| YaL-F11 | Female | Ya-U | 94.10 | 33.73 | 4,849 | 360,358,685 | 17,971 | 21,695 | 17,977 | 93,589 | 11,515 |
| YaL-F13 | Female | Ya-U | 93.45 | 25.51 | 5,686 | 394,307,353 | 19,417 | 23,452 | 18,662 | 89,176 | 11,138 |
| YaL-F20 | Female | Ya-U | 93.54 | 29.50 | 3,974 | 433,524,338 | 17,204 | 19,105 | 23,613 | 73,137 | 5,377 |
| YaL-M7 | Male | Ya-U | 93.53 | 29.53 | 6,035 | 561,508,794 | 28,237 | 41,876 | 17,766 | 51,394 | 55,289 |
| YaL-M8 | Male | Ya-U | 92.27 | 34.70 | 10,432 | 607,100,629 | 33,158 | 63,066 | 13,121 | 41,571 | 86,745 |
| YaL-M10 | Male | Ya-U | 94.12 | 36.13 | 4,390 | 355,606,474 | 16,423 | 19,660 | 19,152 | 92,067 | 4,784 |
| YaL-M11 | Male | Ya-U | 93.65 | 23.32 | 12,479 | 609,683,092 | 33,128 | 61,585 | 13,338 | 41,236 | 68,388 |
| YaL-M14 | Male | Ya-U | 93.87 | 26.68 | nd | 264,551,372 | 18,128 | 18,667 | 14,369 | 104,384 | 3,551 |
| YaL-M16 | Male | Ya-U | 93.98 | 24.02 | 3,289 | 303,933,539 | 19,032 | 20,327 | 15,565 | 97,427 | 4,448 |
| YaL-M17 | Male | Ya-U | 93.68 | 23.79 | 5,858 | 423,920,287 | 20,835 | 25,940 | 18,664 | 82,598 | 23,729 |
| YaL-M18 | Male | Ya-U | 93.78 | 31.66 | 3,193 | 243,577,016 | 17,728 | 19,919 | 12,673 | 105,508 | 5,481 |
| YaL-M21 | Male | Ya-U | 93.34 | 32.54 | 4,996 | 415,438,638 | 18,870 | 23,460 | 19,622 | 81,499 | 6,258 |
| YaL-M23 | Male | Ya-U | 92.95 | 25.93 | 2,040 | 216,002,997 | 18,455 | 19,183 | 11,484 | 111,466 | 3,611 |
| YaL-M25 | Male | Ya-U | 93.89 | 27.16 | 7,512 | 631,952,124 | 33,607 | 48,596 | 16,255 | 36,313 | 57,931 |

Ar = Aripo River, Ya = Yarra River, Qu = Quare River; D = Downstream/High-predation population, U = Upstream/Low-predation population
nd = Supernova could not determine molecule length, likely due to sub-optimal input DNA quality, but the assembly process was successful.

**Supplementary Table S3.** Assembly statistics for the *de-novo* assemblies of the three rivers.

| | Aripo | Quare | Yarra | NCBI genome |
|---|---|---|---|---|
| **# scaffolds** | 11,150 | 8,382 | 15,147 | nd |
| **Longest scaffold (bp)** | 16,773,556 | 22,850,742 | 10,339,082 | nd |
| **Scaffold N50** | 51 | 36 | 121 | nd |
| **Scaffold L50 (bp)** | 3,972,843 | 6,017,975 | 1,509,777 | nd |
| **Scaffold N90** | 334 | 194 | 992 | nd |
| **Scaffold L90 (bp)** | 135,571 | 362,480 | 37,420 | nd |
| **% genome > 50kb** | 92.13 | 93.87 | 89.38 | nd |
| **Total length (bp)*** | 683,635,408 | 688,342,185 | 681,071,722 | 679,931,942 |
| **% gaps (N)*** | 3.29 | 3.89 | 3.45 | 8.45 |
| **BUSCO - Actinopterygii** | | | | |
| **complete** | 96.3 | 96.5 | 96.0 | 95.7 |
| **single-copy** | 94.3 | 94.4 | 93.5 | 93.8 |
| **duplicated** | 2.0 | 2.1 | 2.5 | 1.9 |
| **fragmented** | 1.4 | 1.2 | 1.6 | 1.7 |
| **missing** | 2.3 | 2.3 | 2.4 | 2.6 |

nd – not determined
* calculated on anchored linkage groups only

**Supplementary Table S4.** Distribution of male-linked SNPs by chromosome. SNPs for which all but 3 males (>66%) in each population are heterozygous and all females are homozygous were classified as male-linked.

| | Aripo | | Quare | | Yarra | |
|---|---|---|---|---|---|---|
| **Chromosome** | **Down.** | **Up.** | **Down.** | **Up.** | **Down.** | **Up.** |
| 1 | 87 | 118 | 208 | 56 | 53 | 205 |
| 2 | 118 | 58 | 243 | 62 | 52 | 327 |
| 3 | 81 | 49 | 194 | 32 | 10 | 111 |
| 4 | 68 | 58 | 170 | 45 | 10 | 52 |
| 5 | 54 | 61 | 197 | 46 | 92 | 265 |
| 6 | 84 | 154 | 89 | 99 | 24 | 89 |
| 7 | 122 | 139 | 162 | 80 | 163 | 247 |
| 8 | 35 | 53 | 101 | 604 | 42 | 323 |
| 9 | 94 | 39 | 208 | 76 | 21 | 199 |
| 10 | 132 | 119 | 244 | 85 | 121 | 172 |
| 11 | 46 | 75 | 120 | 28 | 116 | 82 |
| 12* | 283 | 235 | 916 | 1504 | 561 | 4645 |
| 13 | 60 | 77 | 156 | 82 | 85 | 287 |
| 14 | 59 | 60 | 249 | 79 | 17 | 197 |
| 15 | 72 | 51 | 143 | 58 | 207 | 170 |
| 16 | 71 | 55 | 175 | 55 | 24 | 168 |
| 17 | 149 | 79 | 135 | 59 | 93 | 67 |
| 18 | 23 | 38 | 89 | 64 | 42 | 137 |
| 19 | 28 | 54 | 155 | 75 | 203 | 98 |
| 20 | 50 | 45 | 120 | 91 | 98 | 297 |
| 21 | 67 | 91 | 135 | 54 | 411 | 104 |
| 22 | 152 | 61 | 93 | 32 | 26 | 144 |
| 23 | 48 | 73 | 86 | 97 | 33 | 207 |

Down. – downstream population
Up. – upstream population
* sex chromosome

**Supplementary Table S5.** Average nucleotide diversity by population. Diversity was estimated as the proportion of segregating sites (Watterson's theta) in 50 kb windows and averaged across all windows.

| | | Genome | Sex chromosome |
|---|---|---|---|
| **Aripo** | **Downstream** | 0.00360 | 0.00485 |
| | **Upstream** | 0.00237 | 0.00251 |
| **Quare** | **Downstream** | 0.00305 | 0.00387 |
| | **Upstream** | 0.00090 | 0.00163 |
| **Yarra** | **Downstream** | 0.00208 | 0.00287 |
| | **Upstream** | 0.00220 | 0.00234 |

**Supplementary Table S6**. Identified genes on putative Y-linked scaffolds. The total number of annotated genes was 196. Sixty proteins of unknown function are not represented in the table (this includes the 3 proteins of unknown function found in the Aripo watershed).

| Gene name | Description | Count | River* | Genbank** | Chrom*** |
|---|---|---|---|---|---|
| ADD3 | Gamma-adducin | 2 | Ya | XM_017305457 | 1 |
| adra1a | Alpha-1A adrenergic receptor | 1 | Qu | XM_008424673 | 12 |
| AKNA | Microtubule organization protein AKNA | 1 | Qu | XM_008424637 | 12 |
| ALAD | Delta-aminolevulinic acid dehydratase | 1 | Qu | XM_008424693 | 12 |
| ARHGAP11A | Rho GTPase-activating protein 11A | 1 | Qu | XM_008397378 | 21 |
| ARRDC1 | Arrestin domain-containing protein 1 | 2 | Qu | XM_008424643 | 12 |
| Atp8b2 | Phospholipid-transporting ATPase ID | 2 | Ya,Qu | XM_008424732 | 12 |
| Aven | Cell death regulator Aven | 1 | Qu | XM_017302122 | 21 |
| BARHL1 | BarH-like 1 homeobox protein | 1 | Qu | XM_008424665 | 12 |
| BNIP3L | BCL2/adenovirus E1B 19 kDa protein-interacting protein 3-like | 1 | Qu | XM_008424670 | 12 |
| brcc3 | Lys-63-specific deubiquitinase BRCC36 | 1 | Qu | XM_008424562 | 12 |
| C6 | Complement component C6 | 2 | Ya,Qu | XM_017307761 | 12 |
| C7 | Complement component C7 | 2 | Ya,Qu | XM_008403687 | unplaced |
| CACNA1A | Voltage-dependent P/Q-type calcium channel subunit alpha-1A | 1 | Qu | XM_017307837 | 12 |
| Cacna1b | Voltage-dependent N-type calcium channel subunit alpha-1B | 1 | Qu | XM_017307828 | 12 |
| CCDC152 | Coiled-coil domain-containing protein 152 | 1 | Ya | XM_008438664 | 20 |
| ccnb1 | G2/mitotic-specific cyclin-B1 | 1 | Qu | XM_008424602 | 12 |
| CCNG2 | Cyclin-G2 | 2 | Ya,Qu | XM_008424569 | 12 |
| CCNI | Cyclin-I | 2 | Ya,Qu | XM_008424570 | 12 |
| CEL | Bile salt-activated lipase | 1 | Qu | XM_017307734 | 12 |
| CFAP44 | Cilia- and flagella-associated protein 44 | 2 | Ya,Qu | XM_008424558 | 12 |
| CGNL1 | Cingulin-like protein 1 | 1 | Qu | XM_008402566 | unplaced |
| Chek2 | Serine/threonine-protein kinase Chk2 | 1 | Ya | XM_017303285 | unplaced |
| CHRM5 | Muscarinic acetylcholine receptor M5 | 1 | Qu | XM_008397381 | 21 |

| | | | | | |
|---|---|---|---|---|---|
| COL1A2 | Collagen alpha-2 | 1 | Qu | XM_017307678 | 12 |
| DAB2IP | Disabled homolog 2-interacting protein | 1 | Ya | XM_008402526 | unplaced |
| daf-36 | Cholesterol 7-desaturase | 1 | Qu | XM_008424701 | 12 |
| dnajc25 | DnaJ homolog subfamily C member 25 | 1 | Ya | XM_017307726 | 12 |
| DNM1 | Dynamin-1 | 1 | Ya | XM_017303277 | 12 |
| DPYSL2 | Dihydropyrimidinase-related protein 2 | 1 | Qu | XM_017307911 | 12 |
| efna5b | Ephrin-A5b | 1 | Qu | XM_008424718 | 12 |
| Ehmt1 | Histone-lysine N-methyltransferase EHMT1 | 1 | Qu | XM_008424610 | 12 |
| ENTPD2 | Ectonucleoside triphosphate diphosphohydrolase 2 | 1 | Qu | XM_017307733 | 12 |
| entr1 | Endosome-associated-trafficking regulator 1 | 1 | Ya | XM_008402532 | unplaced |
| fam160b2 | Protein FAM160B2 | 2 | Ya | XM_008402518 | unplaced |
| fech | Ferrochelatase, mitochondrial | 2 | Qu | XM_008424594 | 12 |
| FIBCD1 | Fibrinogen C domain-containing protein 1 | 1 | Ya | XM_017303287 | unplaced |
| Fmn1 | Formin-1 | 1 | Qu | XM_008397379 | 21 |
| G2E3 | G2/M phase-specific E3 ubiquitin-protein ligase | 1 | Qu | XM_008424863 | 13 |
| GFI1B | Zinc finger protein Gfi-1b | 1 | Qu | XM_008424660 | 12 |
| GHR | Growth hormone receptor | 2 | Ya,Qu | XM_017307740 | 12 |
| GLUL | Glutamine synthetase | 1 | Qu | | |
| GRIN1 | Glutamate receptor ionotropic, NMDA 1 | 1 | Qu | XM_017307813 | 12 |
| GSN | Gelsolin | 2 | Ya,Qu | XM_017307920 | 12 |
| GSTT1 | Glutathione S-transferase theta-1 | 1 | Ya | XM_008402531 | unplaced |
| GTF3C4 | General transcription factor 3C polypeptide 4 | 1 | Qu | XM_008424662 | 12 |
| GTF3C5 | General transcription factor 3C polypeptide 5 | 1 | Qu | XM_008424702 | 12 |
| htr1a-B | 5-hydroxytryptamine receptor 1A-beta | 2 | Ya,Qu | XM_008424748 | 12 |
| IGSF9B | Protein turtle homolog B | 1 | Qu | XM_017307737 | 12 |
| KCNV2 | Potassium voltage-gated channel subfamily V member 2 | 2 | Ya,Qu | XM_008424587 | 12 |
| Klk1b11 | Kallikrein 1-related peptidase b11 | 1 | Qu | XM_008424614 | 12 |
| MAMDC4 | Apical endosomal glycoprotein | 1 | Qu | XM_017307732 | 12 |

| | | | | | |
|---|---|---|---|---|---|
| Man1b1 | Endoplasmic reticulum mannosyl-oligosaccharide 1,2-alpha-mannosidase | 1 | Qu | XM_008424628 | 12 |
| MFSD14A | Hippocampus abundant transcript 1 protein | 2 | Ya,Qu | XM_017307779 | 12 |
| 00000088 | Similar to Actin, alpha cardiac | 1 | Qu | XM_008397375 | 21 |
| 00000094 | Similar to Ryanodine receptor 3 | 1 | Qu | | |
| NARS1 | Asparagine--tRNA ligase, cytoplasmic | 2 | Ya,Qu | XM_008424576 | 12 |
| Niban2 | Protein Niban 2 | 1 | Qu | XM_008424642 | 12 |
| Nim1k | Serine/threonine-protein kinase NIM1 | 2 | Ya,Qu | XM_008424736 | 12 |
| npr2 | Atrial natriuretic peptide receptor 2 | 2 | Qu | XM_017307730 | 12 |
| OARD1 | ADP-ribose glycohydrolase OARD1 | 2 | Qu | XM_017303228 | unplaced |
| ONECUT2 | One cut domain family member 2 | 1 | Ya | XM_008424565 | 12 |
| Os07g0515000 | tRNA wybutosine-synthesizing protein 2/3/4 | 1 | Qu | XM_017307777 | 12 |
| Parm1 | Prostate androgen-regulated mucin-like protein 1 homolog | 1 | Qu | XM_008424751 | 12 |
| PHPT1 | 14 kDa phosphohistidine phosphatase | 1 | Qu | | |
| PIGO | GPI ethanolamine phosphate transferase 3 | 1 | Qu | XM_008424720 | 12 |
| PPP2R2A | Serine/threonine-protein phosphatase 2A 55 kDa regulatory subunit B alpha isoform | 1 | Qu | XM_017307735 | 12 |
| pum3 | Pumilio homolog 3 | 2 | Ya,Qu | XM_017307725 | 12 |
| RAB27B | Ras-related protein Rab-27B | 3 | Ya,Qu | XM_008424581 | 12 |
| RAPGEF1 | Rap guanine nucleotide exchange factor 1 | 1 | Ya | XM_017303277 | unplaced |
| rasgrf2 | Ras-specific guanine nucleotide-releasing factor 2 | 1 | Qu | XM_008423269 | 12 |
| RYR3 | Ryanodine receptor 3 | 2 | Qu | XM_017302123 | 21 |
| Scg5 | Neuroendocrine protein 7B2 | 1 | Qu | | |
| Sh3bp2 | SH3 domain-binding protein 2 | 2 | Ya,Qu | XM_008403810 | unplaced |
| SHROOM3 | Protein Shroom3 | 4 | Ya,Qu | XM_017307728 | 12 |
| SOWAHB | Ankyrin repeat domain-containing protein SOWAHB | 2 | Ya,Qu | XM_008424573 | 12 |
| SPAG8 | Sperm-associated antigen 8 | 1 | Qu | XM_017307982 | 12 |
| SPIN1 | Spindlin-1 | 1 | Ya | XM_017306593 | 9 |

| | | | | | |
|---|---|---|---|---|---|
| SPINW | Spindlin-W | 1 | Ya | | |
| SPINZ | Spindlin-Z | 1 | Qu | | |
| Ssna1 | Sjoegren syndrome nuclear autoantigen 1 homolog | 1 | Qu | XM_008424608 | 12 |
| ST8SIA3 | Sia-alpha-2,3-Gal-beta-1,4-GlcNAc-R | 1 | Ya | XM_008424566 | 12 |
| STOM | Erythrocyte band 7 integral membrane protein | 1 | Qu | XM_017307919 | 12 |
| Stoml2 | Stomatin-like protein 2, mitochondrial | 1 | Qu | XM_008424721 | 12 |
| Stxbp1 | Syntaxin-binding protein 1 | 1 | Qu | XM_008424639 | 12 |
| Taf1c | TATA box-binding protein-associated factor, RNA polymerase I, subunit C | 1 | Qu | XM_008424568 | 12 |
| Tbx3 | T-box transcription factor TBX3 | 1 | Qu | XM_008403160 | unplaced |
| TCF12 | Transcription factor 12 | 1 | Qu | XM_008402569 | unplaced |
| tmc2a | Transmembrane channel-like protein 2-A | 1 | Qu | | |
| TMEM230 | Transmembrane protein 230 | 1 | Qu | XM_008424674 | 12 |
| TRIM25 | E3 ubiquitin/ISG15 ligase TRIM25 | 1 | Qu | XM_008424560 | 12 |
| Trim29 | Tripartite motif-containing protein 29 | 3 | Ya,Qu | XM_008434801 | 17 |
| TSC1 | Hamartin | 1 | Qu | XM_008424661 | 12 |
| Txnl1 | Thioredoxin-like protein 1 | 3 | Ya,Qu | XM_008424578 | 12 |
| UNC13B | Protein unc-13 homolog B | 2 | Qu | XM_017307753 | 12 |
| Unc5d | Netrin receptor UNC5D | 1 | Qu | XM_017307716 | 12 |
| VLDLR | Very low-density lipoprotein receptor | 2 | Ya,Qu | XM_008424754 | 12 |
| Wdr7 | WD repeat-containing protein 7 | 2 | Ya,Qu | XM_017307727 | 12 |
| Whrn | Whirlin | 2 | Qu | XM_008424630 | 12 |
| ZFAND5 | AN1-type zinc finger protein 5 | 1 | Qu | XM_008424601 | 12 |

* Qu: Quare; Ya: Yarra

** Genbank accession number of the best reciprocal BLAST hit to the NCBI reference genome annotation

*** Chromosome of the NCBI reference genome

**Supplementary Figure S1. Inversion on the sex chromosome (Chromosome 12) between the reference genome (Künstner et al. 2016) and our female genome assemblies. a)** Dot-plots of Chromosome 12 alignments between the reference genome and our three female assemblies. Forward alignments are drawn in blue and reverse alignments are drawn in red. The Aripo sequence shows a clear rearrangement of the first ~10 Mb in the NCBI reference involving an inversion and translocation to the middle of the chromosome. The Quare and Yarra assemblies did not have long enough scaffolds spanning the full length of the inversion, but inverted breakpoints (in red) are clearly visible at the same genomic positions of the NCBI genome as those found in Aripo. **b)** IGV snapshot around the inferred breakpoint positions in the NCBI reference. From read alignments, we identified clear breakpoints in all three female genomes for which the reads surrounding the breakpoints do not have the expected pair orientation (light blue – panel b)) and/or are soft-clipped at the breakpoint positions (panel c). **c)** Inferred structure of the genomic rearrangement between a target genome and the reference. The inverted segment is shown in red and approximate breakpoint positions are indicated. Arrows indicate the expected read-pair orientation of reads mapping near the breakpoints of both configurations.

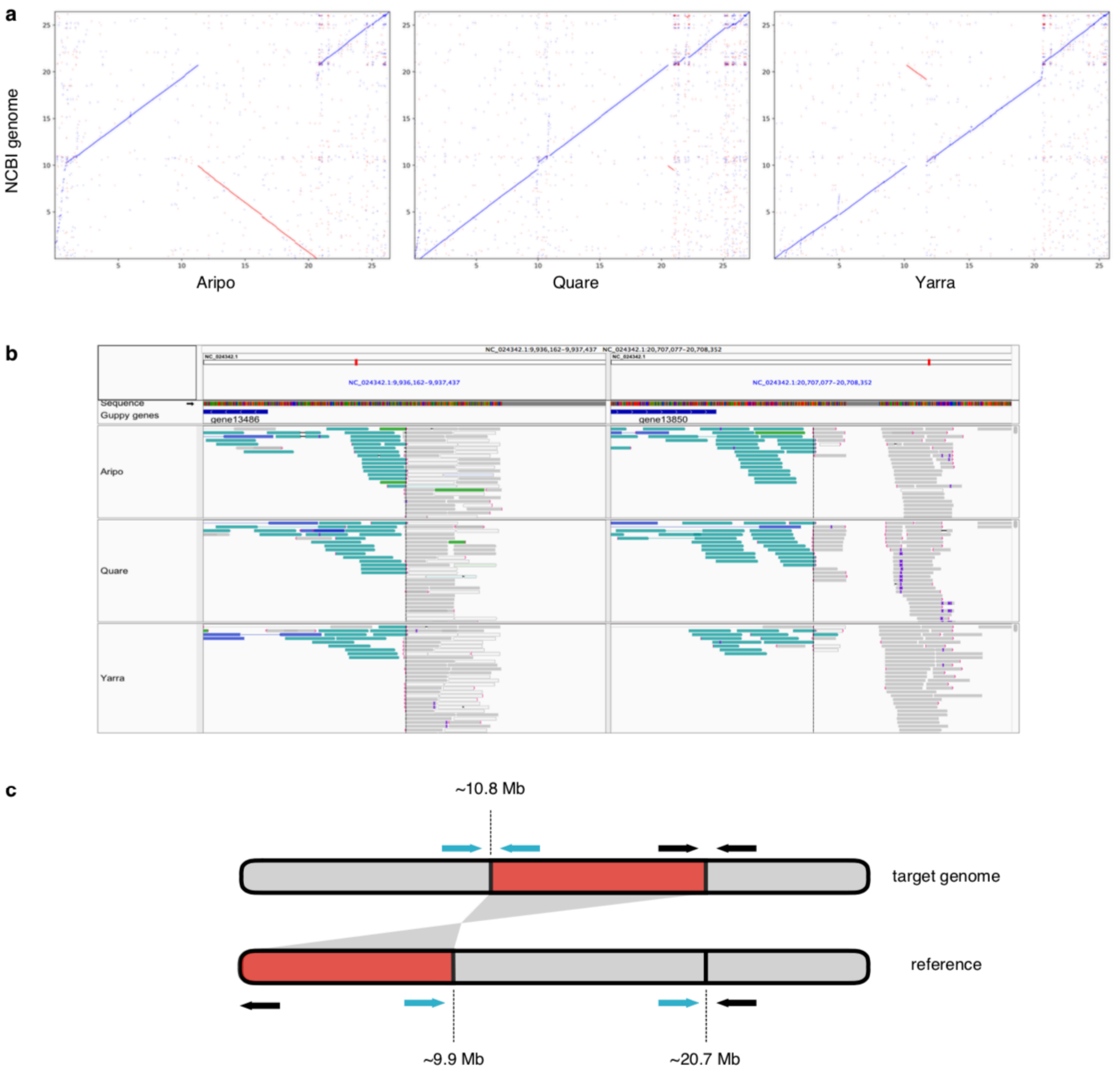

**Supplementary Figure S2. Detection of the ancestral region of recombination suppression in the sex chromosome (Stratum I). a)** Male to female coverage differences in Chromosome 12 for downstream, high-predation (black) and upstream, low-predation (orange) populations in the Aripo (**a**), Quare (**b**) and Yarra (**c**) watersheds. The $\log_2$ of male to female coverage was calculated in non-overlapping windows of 50 kb. The 95% confidence interval, inferred from bootstrapping autosomal regions is shaded in grey. The right-side density plots show the frequency (counts) of windows. ***, p-value < 0.001.

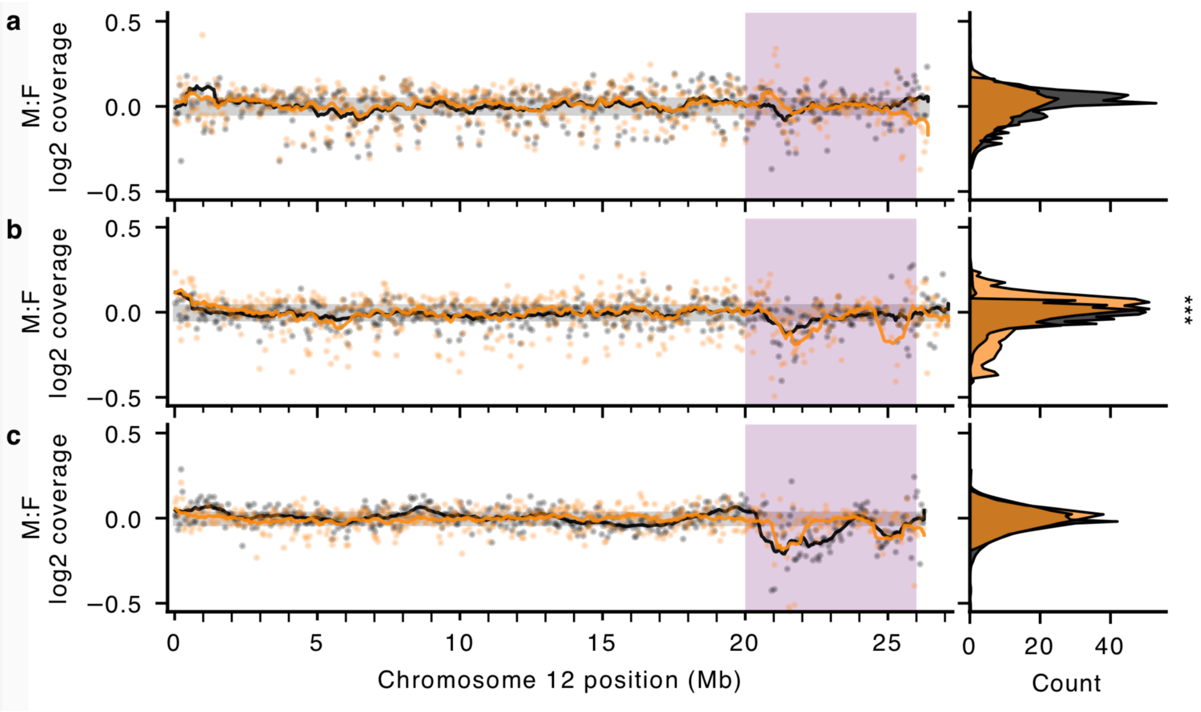

**Supplementary Figure S3. Individual read coverage for Stratum I.** Normalised median read coverage in non-overlapping windows of 50 kb. Coverage is plotted for each individual with males in blue and females in red.

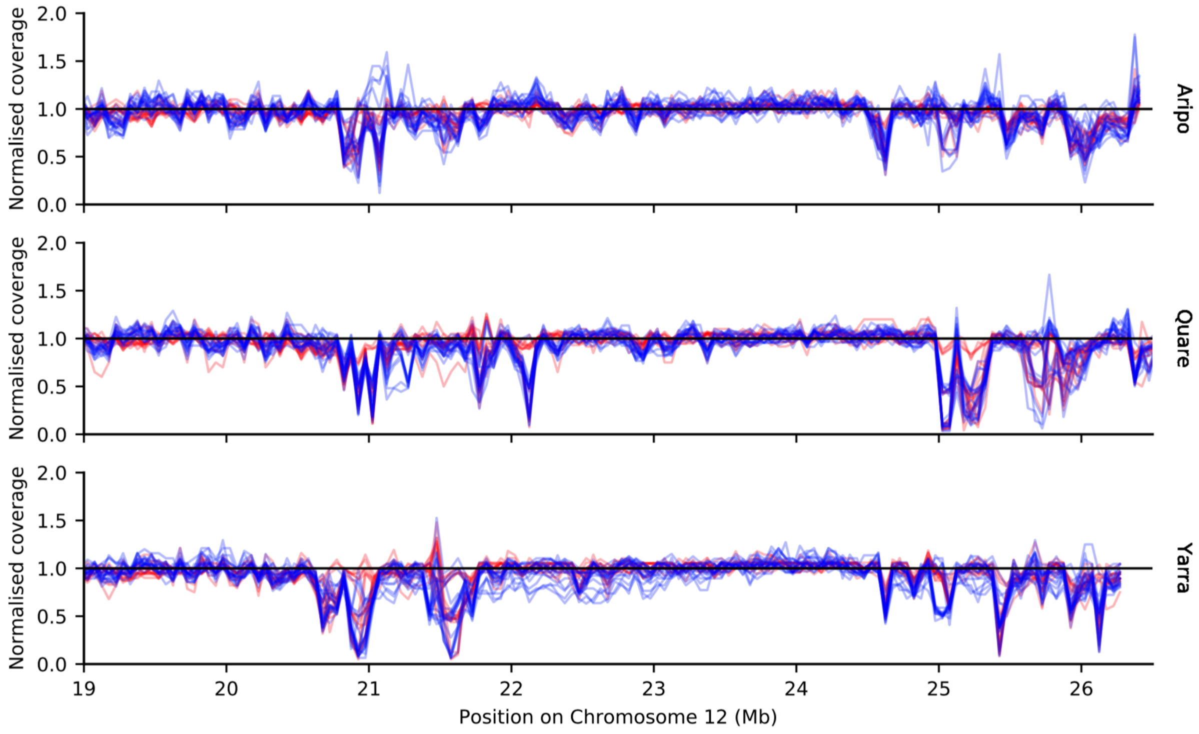

**Figure S4. Evidence of incomplete lineage sorting in the pseudo-autosomal region (PAR) and Stratum I (SDR) of the sex chromosome.** Principal components analysis (PCA) for all samples in the Aripo, Quare and Yarra watersheds. Panel **a** includes all SNPs on Chromosome 12 except for those found within Stratum I. Panel **b** includes all SNPs found within Stratum I. Males are shaded in blue and females are shaded in red.

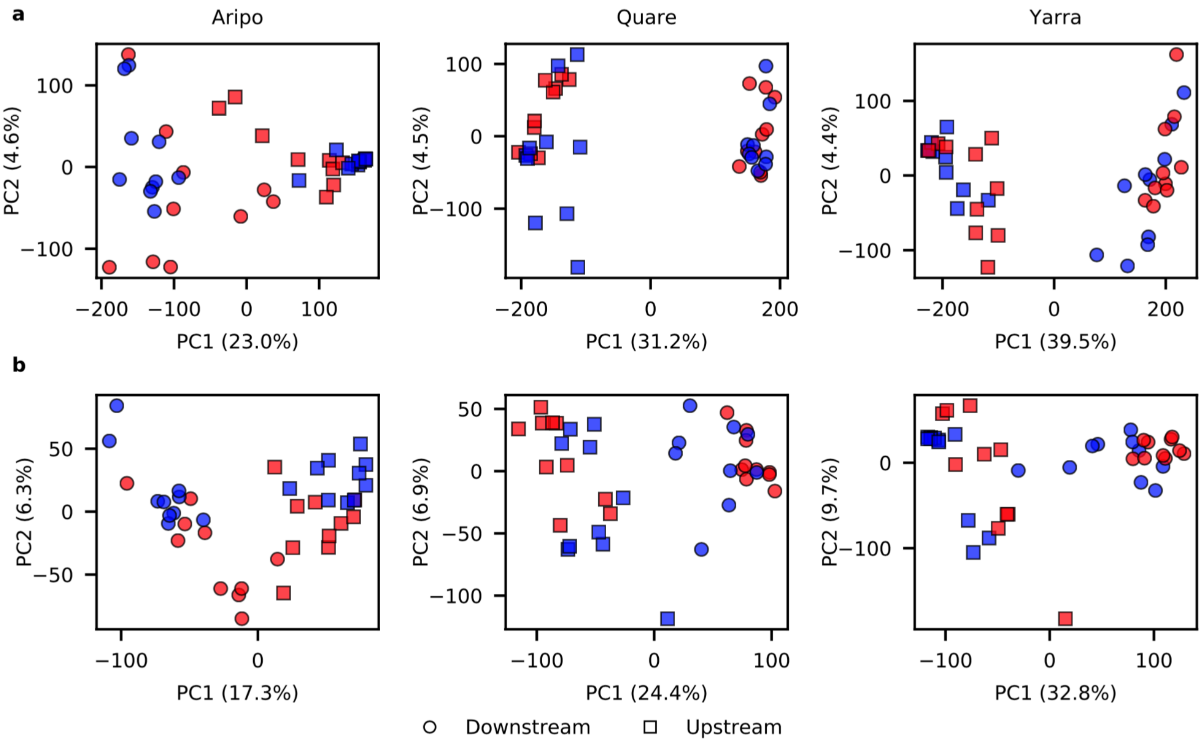

**Figure S5. Long-range barcode overlap in males and females.** 10x Genomics barcode overlap calculated in non-overlapping windows of 20 kb with all samples of each sex combined. The distribution of barcode overlaps is similar between males and females, suggesting that sex-chromosome divergence in the guppy is probably not driven by a large chromosomal rearrangement (e.g. inversion). **a-c)** Male plots in Aripo (a), Quare (b) and Yarra (c). **d-f)** Female plots in Aripo (d), Quare (e) and Yarra (f).

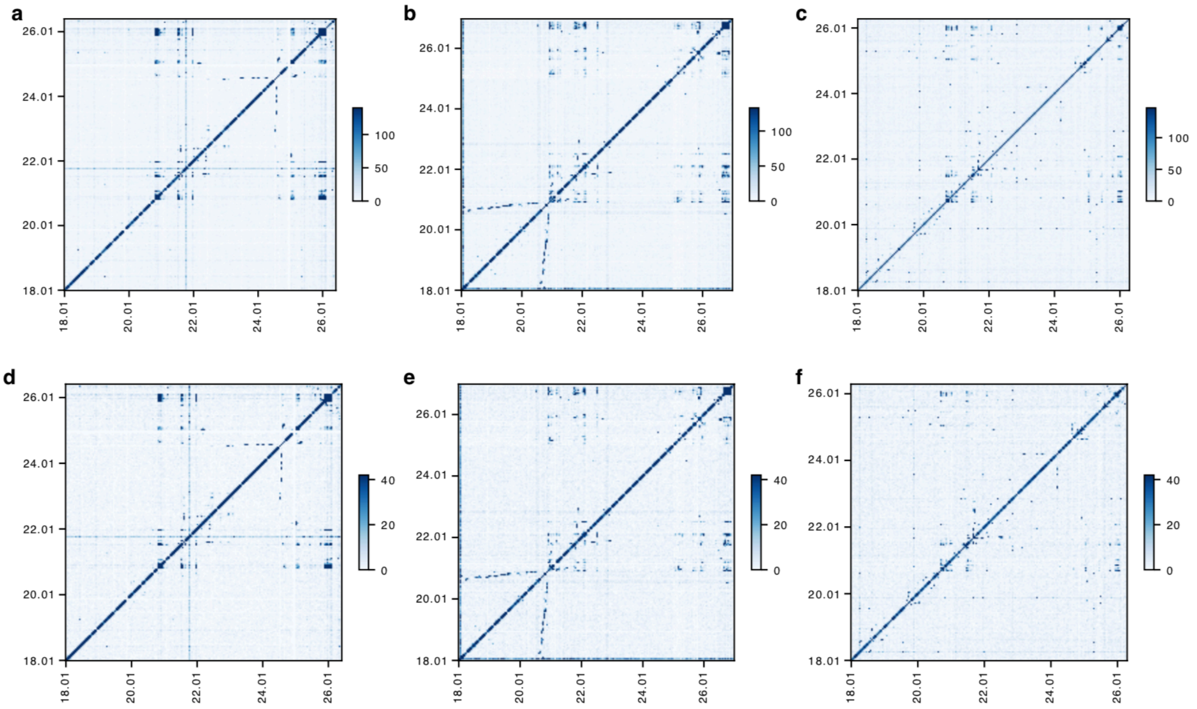

**Supplementary Figure S6. Phase-switch error rate estimated for the samples sequenced with 10x Genomics.** We calculated phase-switching errors after the computational phasing of all populations for the phase sets in each individual originally reconstructed with the 10x Genomics Long Ranger pipeline. The plots show the average phase switching (y axis) against the average phase length (x axis) per sample. Male samples are shaded in blue and females in red. The dotted line indicates a switching error rate of 5%.

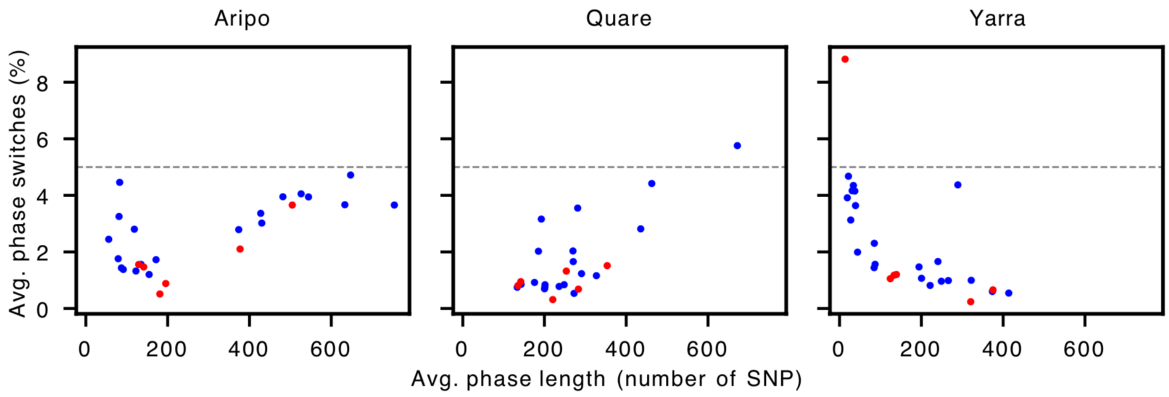

**Supplementary Figure S7. Haplotype networks in the Quare River.** Haplotypes identified in both upstream and downstream populations are shown. The circle area is proportional to haplotype frequency with the smallest circles representing single haplotypes, and branch lengths connecting haplotypes are proportional to the number of SNPs between haplotypes. Haplotypes from downstream, high-predation populations in grey and upstream, low-predation haplotypes in orange. The approximate location (in Mb) of each region on the sex chromosome is indicated above the network.

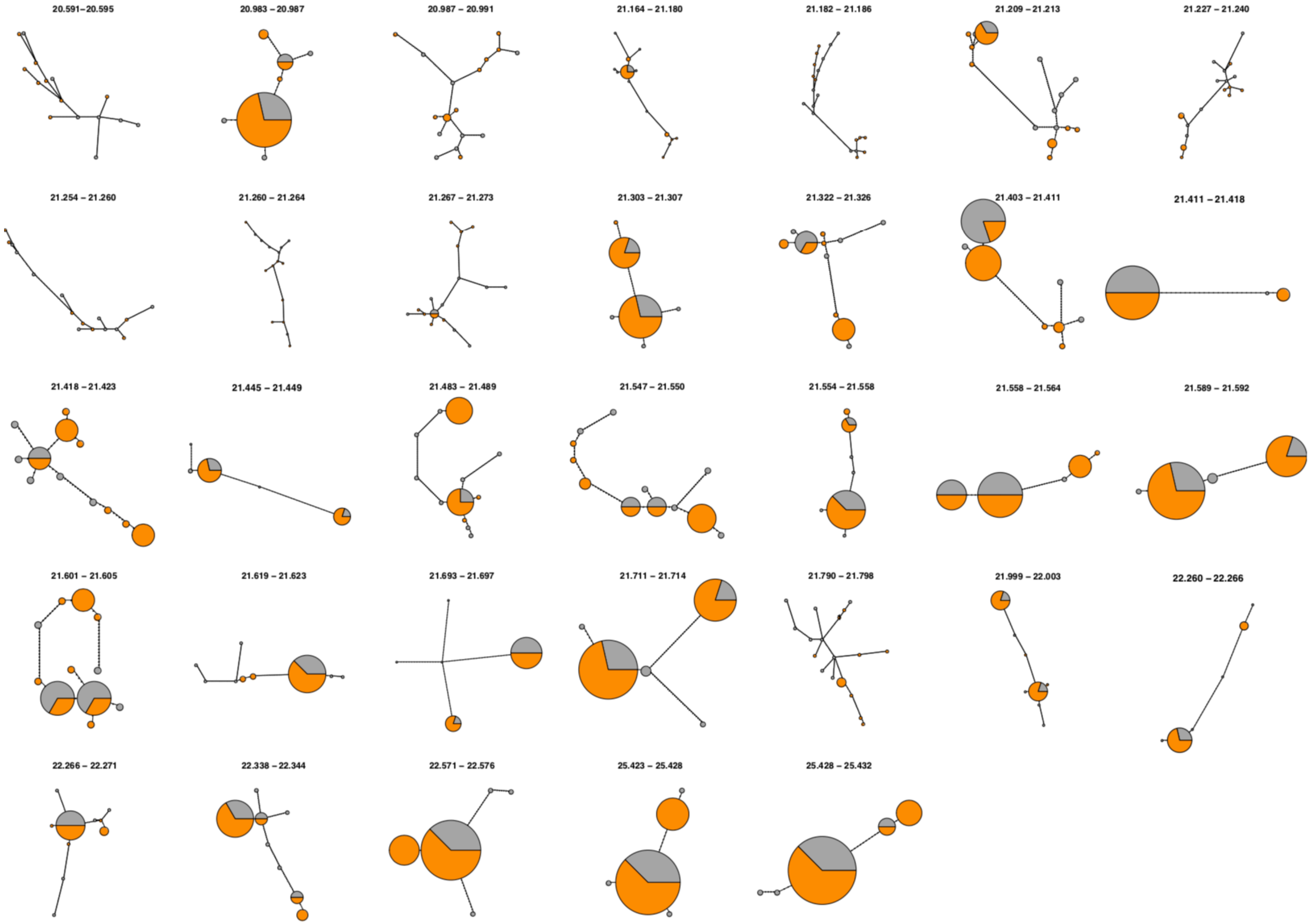

**Supplementary Figure S8. Haplotype networks in the Yarra River.** Haplotypes identified in both upstream and downstream populations are shown. The circle area is proportional to haplotype frequency with the smallest circles representing single haplotypes, and branch lengths connecting haplotypes are proportional to the number of SNPs between haplotypes. Haplotypes from downstream, high-predation populations in grey and upstream, low-predation haplotypes in orange. The approximate location (in Mb) of each region on the sex chromosome is indicated above the network.

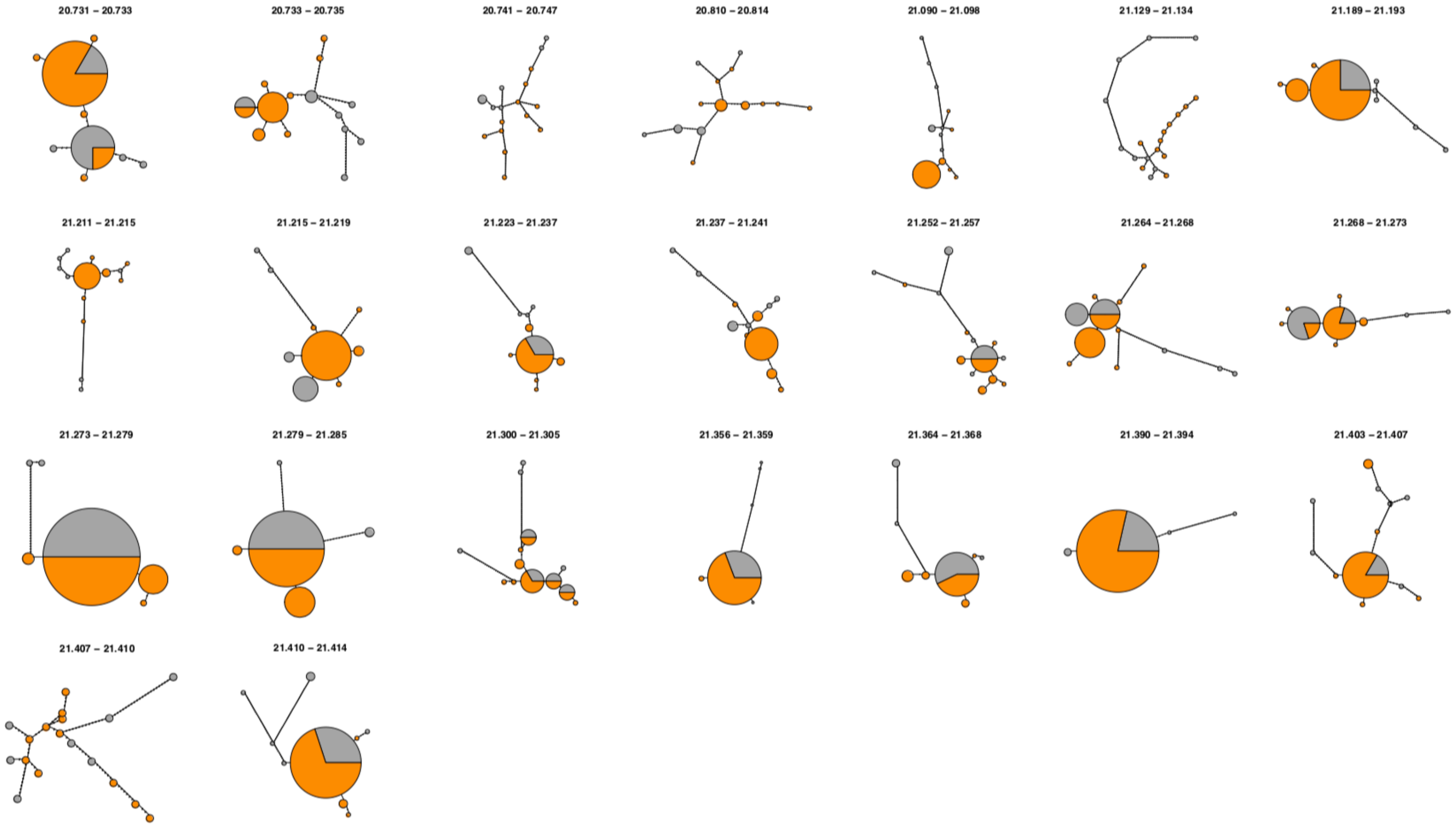

**Supplementary Figure S9. False positive rate (FPR) in Y-mer discovery. a)** Heatmaps of male- (M) and female- (F) specific k-mer counts in all populations. Numbers in the last column indicate the number of sex-specific k-mers fixed in each population. **b)** We used the FPR to estimate the frequency of a k-mer in the male population that can be confidently identified as a Y-mer. First, we recorded the total number of k-mers observed in only one sex at different frequencies in the population, between 1 and 10 individuals (a). Assuming that female-specific k-mers are false positives, the FPR of Y-mers was then estimated, at each frequency, as the number of female-specific k-mers relative to the number of male-specific k-mers minus female-specific k-mers. The dashed horizontal line indicates an FPR of 5%. Note that the slight increase in the FPR in the Quare-upstream population is lower than 5% (0.048).

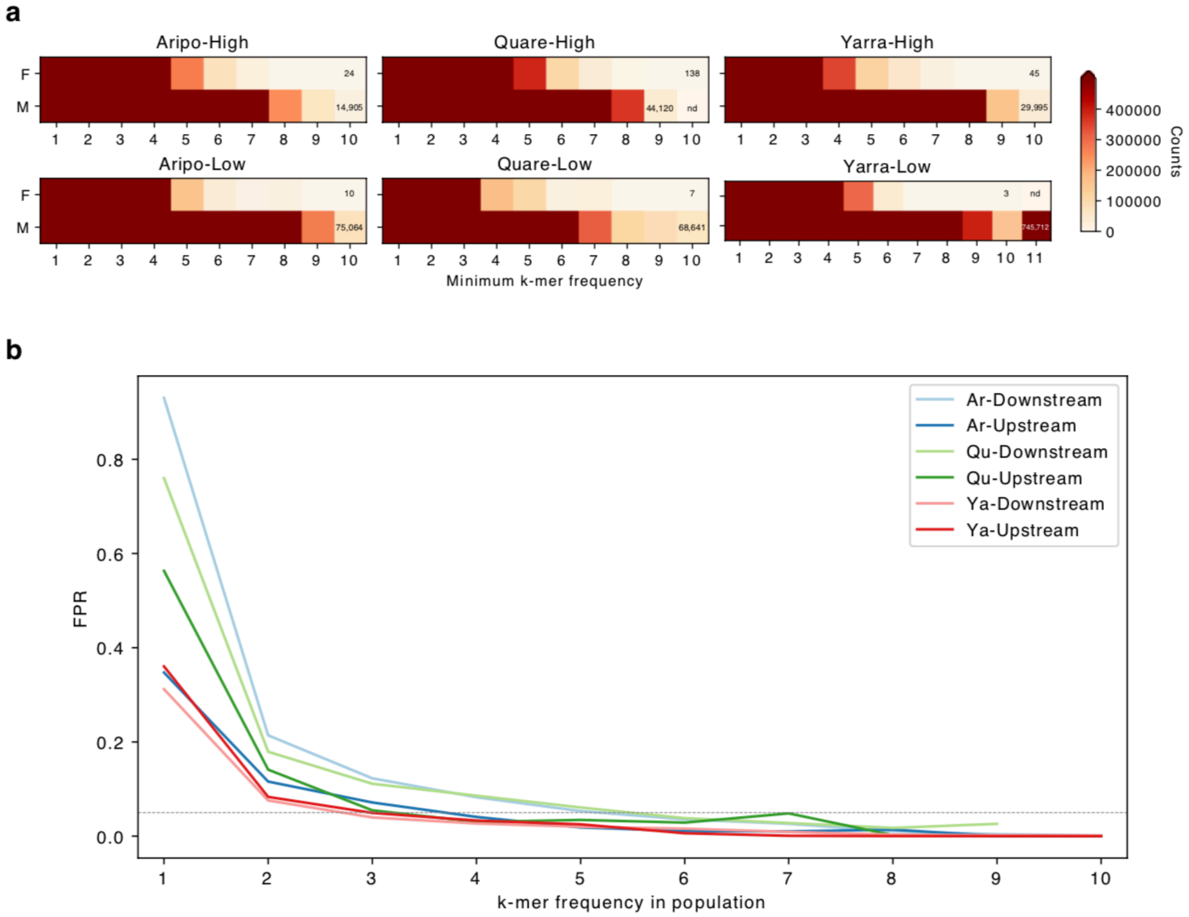

**Supplementary Figure S10. Most Y-mers align non-uniquely to the genome.** The barplots show the total percentage of Y-mers with unique and non-unique (repetitive) alignments to the respective reference genomes of each watershed. Given that Y-mers are male-specific by definition and the reference genomes come from females, we allowed for 1 difference, mismatch or gap, between Y-mers and the reference genome.

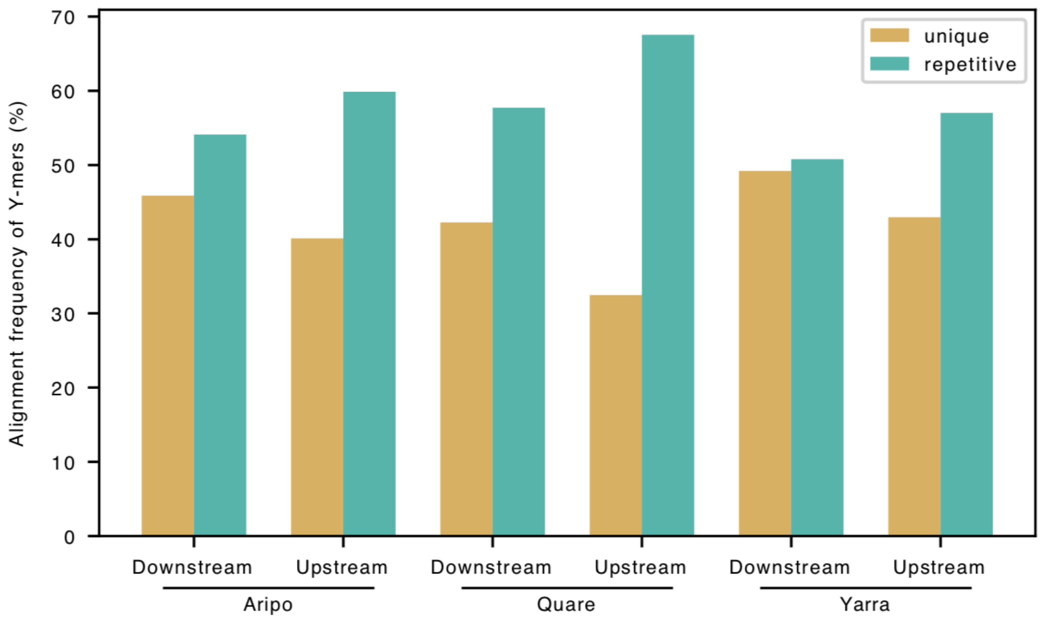

**Supplementary Figure S11. Distribution of transposable elements (TEs) in the guppy sex chromosome.** Boxplots showing the density (total sequence of TEs in every non-overlapping 50 kb window) of all classes of TEs (**a**), and for DNA transposons and Long Interspersed Nuclear Elements (LINEs) (**b**). Statistics were calculated for the Y-linked scaffolds (Y), the region of the X chromosome homologous to the SDR (X) and to the pseudo-autosomal region (PAR). ***p-value < 0.001, **p-value < 0.01, *p-value < 0.05.

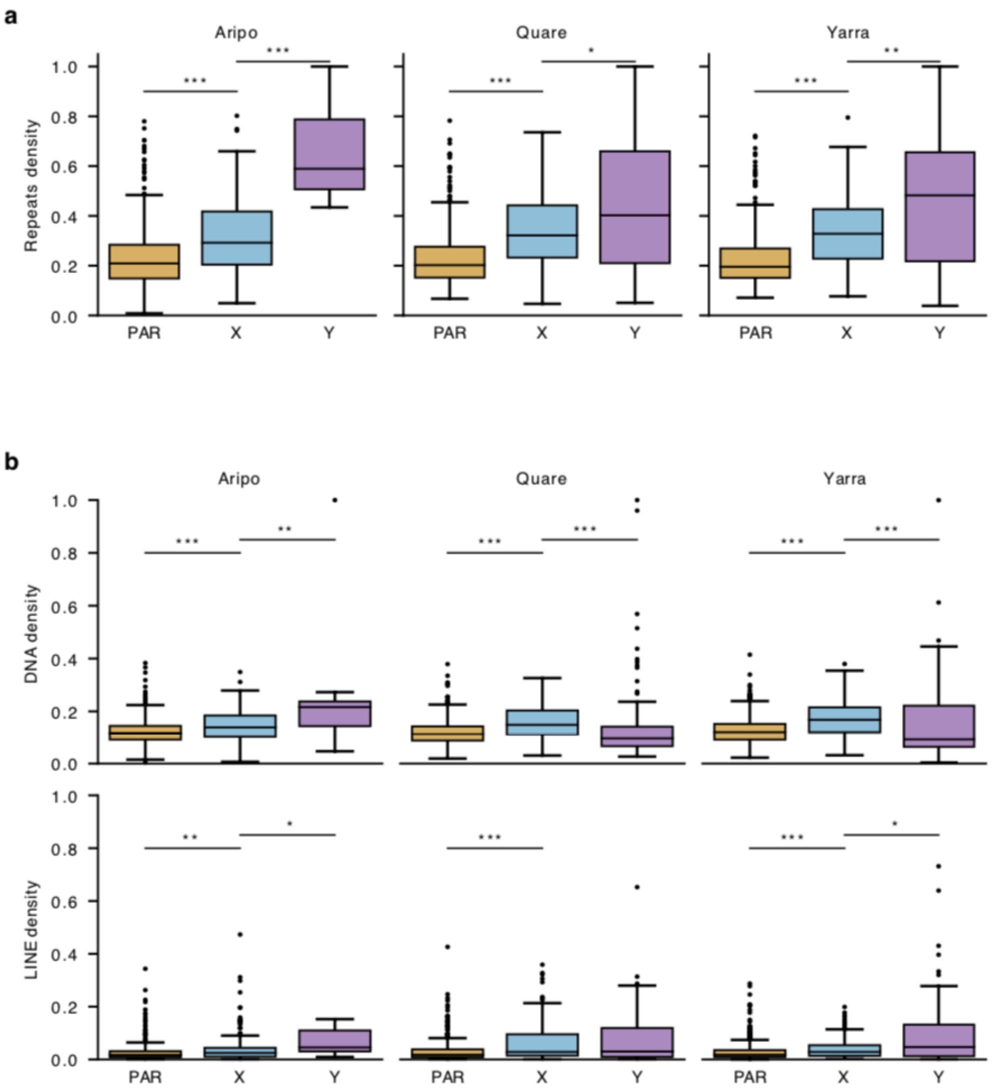
